# In Search of Transcriptomic Correlates of Neuronal Firing-Rate Adaptation across Subtypes, Regions and Species: A Patch-seq Analysis

**DOI:** 10.1101/2024.12.05.627057

**Authors:** John Hongyu Meng, Yijie Kang, Alan Lai, Michael Feyerabend, Wataru Inoue, Julio Martinez-Trujillo, Bernardo Rudy, Xiao-Jing Wang

**Affiliations:** Center for Neural Science, New York University, New York, NY 10003, USA; Department of Physiology and Pharmacology, Western University, London, ON N6A 3K7, Canada; Robarts Research Institute, Western University, London, ON N6A 3K7, Canada; Schulich School of Medicine and Dentistry, Western University, London, ON N6A 3K7, Canada; Neuroscience Institute, NYU Grossman School of Medicine, New York, NY 10016, USA; Department of Neuroscience and Physiology, NYU Grossman School of Medicine, New York, NY 10016, USA; Department of Anesthesiology, Perioperative Care and Pain Medicine, NYU Grossman School of Medicine, New York, NY 10016, USA

## Abstract

Can the transcriptomic profile of a neuron predict its physiological properties? Using a Patch-seq dataset of the primary visual cortex, we addressed this question by focusing on spike rate adaptation (SRA), a well-known phenomenon that depends on small conductance calcium (Ca)-dependent potassium (SK) channels. We first show that in parvalbumin-expressing (PV) and somatostatin-expressing (SST) interneurons, expression levels of genes encoding the ion channels underlying action potential generation are correlated with the half-width (HW) of spikes. Surprisingly, the SK encoding genes are not correlated with the degree of SRA (dAdap). Instead, genes that encode proteins upstream from the SK current, as well as the low-voltage-activated Calcium channel, are correlated with dAdap. This general principle is supported by datasets from the mouse primary motor cortex and multiple macaque brain regions. Finally, we generate testable predictions by constructing a minimal model.

## Introduction

With the advances in the brain connectome and the simultaneous recording from multiple brain areas in behaving animals, studies of the large-scale multiregional brain have come to the fore^1^. To understand brain-wide dynamics, it is necessary to consider differences in the electrophysiological features of single neurons in different areas. However, single-cell physiological studies such as patch-clamp measurements are costly; at the present time there is a dearth of systematic comparisons of single-cell physiological characteristics across many brain regions. On the other hand, single-cell transcriptomic analysis has become common and offers a novel approach to quantification of brain diversity and heterogeneity^2–4^. So far, this tool has been mostly deployed for cell-type classification. Can a transcriptomic profile predict the physiological properties of single cells? The answer is not obvious, as there are multiple intermediate steps from mRNAs to proteins to physiological functions of receptors and ion channels. In this work, we tackled this challenging question by focusing on single-neuron spike rate adaptation, a salient characteristic of firing patterns in distinct subtypes of neurons.

The development of Patch-seq marks a significant advancement in correlating transcriptomic data with single-cell electrophysiological features^5^. Unlike quantitative real-time PCR (RT-PCR), which is limited to studying specific genes with high accuracy^6^, and single-cell RNA sequencing (scRNA-seq), which provides a genome-wide expression profile across diverse neuron types and species^4^, Patch-seq combines scRNA-seq with patch-clamp recordings. This method simultaneously captures RNA sequencing, electrophysiological, and morphological data from the same neuron, providing data at an unprecedented resolution. This offers a novel approach to understanding neural computation at the molecular level. Previous work has performed unsupervised analysis of Patch-seq datasets to generate a list of genes that correlate with electrophysiological or morphological properties^7^. However, this approach typically limits the analysis to correlations between individual genes and features of interest, making it less likely to capture the complex interactions between various genes and provide a causal explanation of the underlying mechanisms.

To accurately predict electrophysiological features from transcriptomic data without overfitting to region-specific genes, we adapt a comprehensive approach that considers the broader network of gene interactions in addition to gene-feature correlations. We started with one important feature of single neurons: spike rate adaptation (SRA)^8^. This feature was first discovered in^9^, describing how neurons modulate their output to the same input. It is important in various cognitive functions that have been validated by experimental and theoretical works alike: sensory processing from vision^10^, audition^11^, and olfaction^12^; working memory in language processing^13^; perceptual bistability^14^ and multistable perceptions^15^; theta sweep in place cells^16^; precision in spike-timing^17^; and processing of temporally dispersed information^18^. Interestingly, different cell types show large differences in the SRA^19^. Parvalbumin (PV) interneurons (IN), recognized by their signature narrow action potential (AP) waveform, show no or little SRA. Somatostatin (SST) INs, which share the same origins from medial ganglionic eminence (MGE) as PV INs^20^, in general have a broader AP waveform and a stronger adaptation compared to PV. PV and SST are widely distributed in the layers and regions of the cortex and account for about 70% of the total INs of the whole cortex^19^, although some variability between cortical areas has been reported^21^. These features make them ideal first targets for studying the mechanisms behind the SRA.

In prior studies, the mechanisms underlying firing-rate adaptation have been examined primarily in pyramidal cells^22^. Typically, an AP activates high-voltage-activated (HVA) calcium (Cav) channels, particularly L-type. The resulting *Ca*^2+^ influx through these L-type Cav channels further activates small-conductance calcium-activated potassium (SK) channels, which generate an outward afterhyperpolarization potential (AHP) that slows the regeneration of subsequent APs^23^.

However, little is known about whether this mechanism generalizes to INs. In addition to the SK pathway, other mechanisms may contribute, such as inactivation of low-voltage-activated (LVA) Cav channels, calcium buffering systems^22^, M-type K^+^ channels, and the hyperpolarization-activated cation (H) current^24^.

Here, we utilize the open Patch-seq dataset^25^ recorded from the primary visual cortex (V1), where both electrophysiological and transcriptomic data are available for PV and SST cells, to search for transcriptomic correlates of firing-rate adaptation. We first show that SRA, measured by the degree of adaptation (dAdap), is a key feature in distinguishing PV and SST cells. We further find that the AP half-width (HW) and dAdap are highly correlated at the single-cell level. Next, we seek the transcriptomic correlation of adaptation using the known transcriptomic correlation of HW as a control. We develop a method to systematically de-noise the original Patch-seq data at the transcriptomic-defined subtype (T-type) level and further validate that HW differences correlate with the transcriptomic data at the T-type level. We hypothesize that the SK channel encoding gene should correlate with dAdap, but we surprisingly find that the data does not support this. Instead, the Ca influx-related mechanisms, namely HW and HVA Cav channel encoding genes, can explain the observed dAdap differences. In addition, the inactivation of LVA Cav channel explains a fraction of the observed differences. We predict other genes that are likely involved in modulating firing-rate adaptation. We further extended our analysis to a mouse M1 Patch-seq dataset that includes pyramidal cells^26^, as well as to an electrophysiological and a transcriptomic dataset^3^ covering the same three brain regions in macaque monkeys, thereby confirming the generality of our conclusions. Finally, we built a Hodgkin-Huxley-type model that reproduces the observed experimental data and generates testable pharmacological predictions.

## Results

### Spike Rate Adaptation and Spike Width are Correlated in PV and SST Interneurons

PV and SST INs together account for about 70% of the total INs of the whole cortex^19^. Here, we analyze the electrophysiological features of PV and SST cells from the open Patch-seq dataset from the mouse V1^25^. PV cells show little adaptation over a 1s-long square-pulse injection current, while SST cells show strong adaptation. Furthermore, PV cells show the signature narrow AP waveform, while SST cells show a wider AP waveform (Figure 1A).

**Figure 1.**
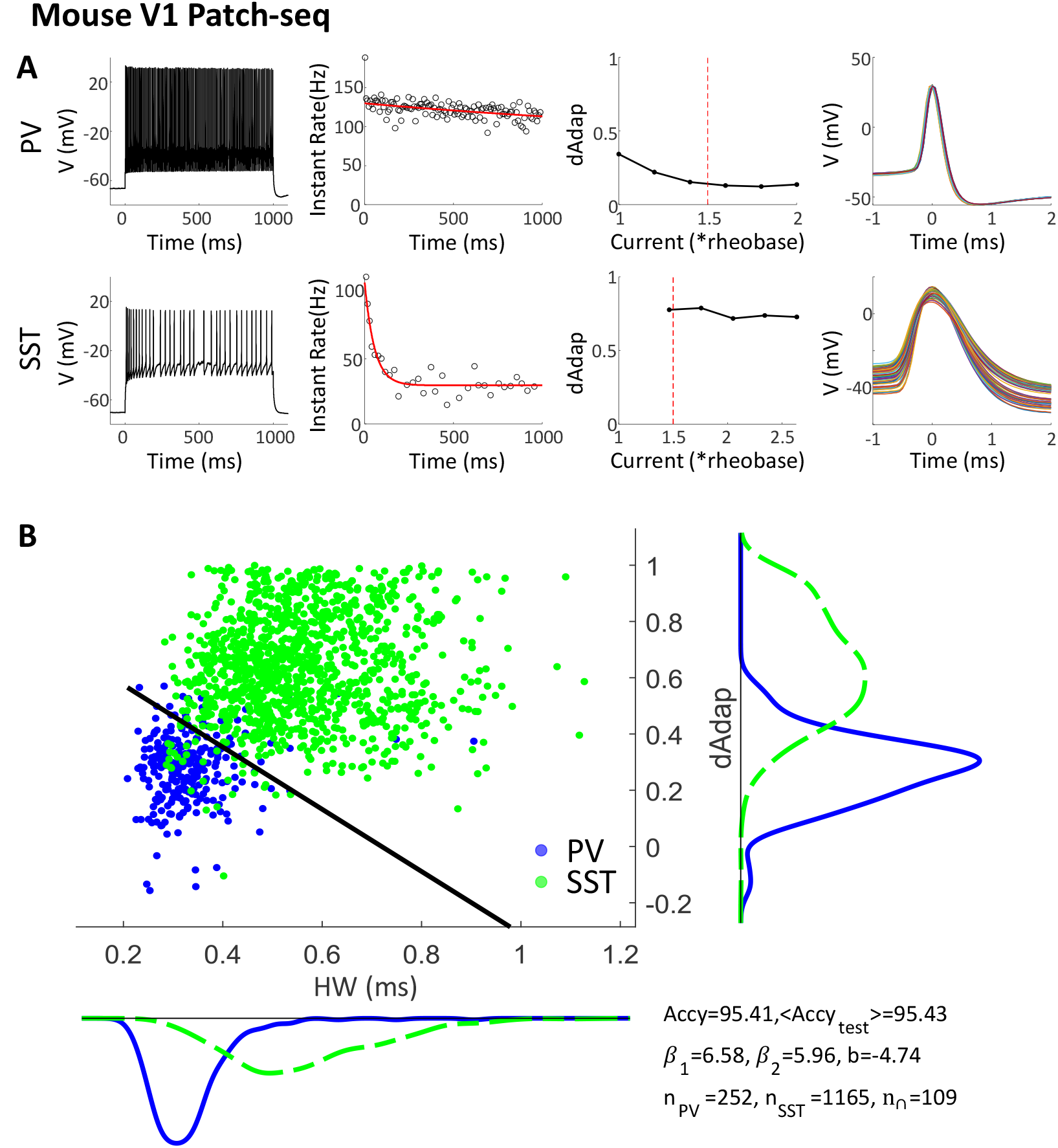
The degree of adaptation and the width of the action potential are highly correlated.(A) Different electrophysiological features of PV and SST INs. From left to right: one recording over a 1s-long square-pulse current injection. Instantaneous firing rate as one over the interspike intervals. The curve is fitted to an exponential decay function. dAdap over injection currents. The red dashed line indicates 1.5× the rheobase. AP shapes are overlaid, while each line represents an AP from the PV cells. (Top) Results from an example PV cell; Bottom, an SST cell. (B) PV and SST cells can be classified based on HW and dAdap. The dashed line is the classification boundary trained by a Support Vector Machine, which is indicated by *y* =−(*xβ*_1_ + *b*)*/β*_2_. See Supp. Spreadsheet 1 for details. Accy, classification accuracy, tested accuracy < Accy_*test*_ > is calculated by the average performance if trained on a randomly split 80% training dataset and tested on the remaining 20% testing set. The sample sizes of PV cells, SST cells, and both *Pvalb*+/*Sst* + cells (see the main text) are indicated by *n*_PV_, *n*_SST_, and *n*_∩_.

We quantitatively measured the firing-rate adaptation by degree of adaptation (dAdap). To do so, we first fit the curve of the instantaneous firing rate to an exponential decay function *f* (*t*) = *a* + *b* exp(−*ct*). After fitting, we compare the firing rate at the end of the 1*s* simulation to the initial firing rate dAdap = 1 − *f* (1)*/f* (0), which varies little at high injection current (above 1.5× rheobase, Figure 1A, 3rd column). In the following analysis, we use the closest sweep above 1.5× rheobase to measure the dAdap for each cell. We quantify the AP waveform by HW, the average time above half of the peak, and the firing threshold for each AP (see Methods).

Next, we investigate how effectively HW and dAdap can distinguish PV and SST cells, compared to other classification methodologies. In^25^, PV and SST cell types are defined through a multi-gene classification algorithm. Based on HW and dAdap along, a trained classifier achieves an accuracy of 95.3% (Figure 1B). Conventionally, classification is performed using marker genes, such as *Pvalb* and *Sst*, like through immunohistochemistry labeling. To compare the effectiveness of multi-gene classification and single marker gene classification, we analyzed the marker gene distribution in the V1 dataset (Supp. Figure 1A). Assuming a threshold at the 20th percentile of *Pvalb* and *Sst* counts per million (CPM) distribution to determine whether a cell is *Pvalb*+ or *Sst* +, respectively, we identified a significant population of *Pvalb*+/*Sst* + cells, resulting in an 89.6% agreement with the multi-gene classification (Supp. Figure 1A). This suggests that a classifier based on HW and dAdap is better aligned with the multi-gene classification than one based on the conventional marker genes. In addition, we did not find a significant difference in classification accuracy when considering laminar differences (Supp. Figure 1B). Moreover, our trained classifier from V1 achieves a 86.4% accuracy on a PV/SST dataset collected from L2/3 of the somatosensory area in mice (Supp. Figure 1C).

### HW and dAdap differences are reflected in the transcriptomic data

Since HW and dAdap differ significantly between PV and SST populations, we next investigated the underlying mechanisms and whether they are reflected in transcriptomic data. We began by performing quality control (Figure 2A, B), including only cells with high total raw counts and numbers of unique detected genes. Additionally, Patch-seq data may include contamination from nearby microglia^27^. Following the approach in^27^, we computed a contamination score for each cell and excluded those with high scores (Figure 2B; see Methods). To ensure reliable dAdap measurements, we also excluded cells not driven to at least 1.5 times rheobase. Threshold variation within a reasonable range yielded qualitatively similar results (Supp. Figure 2). After all quality control steps, approximately 35% of cells remained. We further excluded genes with low expression (see Methods). In subsequent analyses, genes within a family of interest may be omitted if they did not pass quality control.

**Figure 2.**
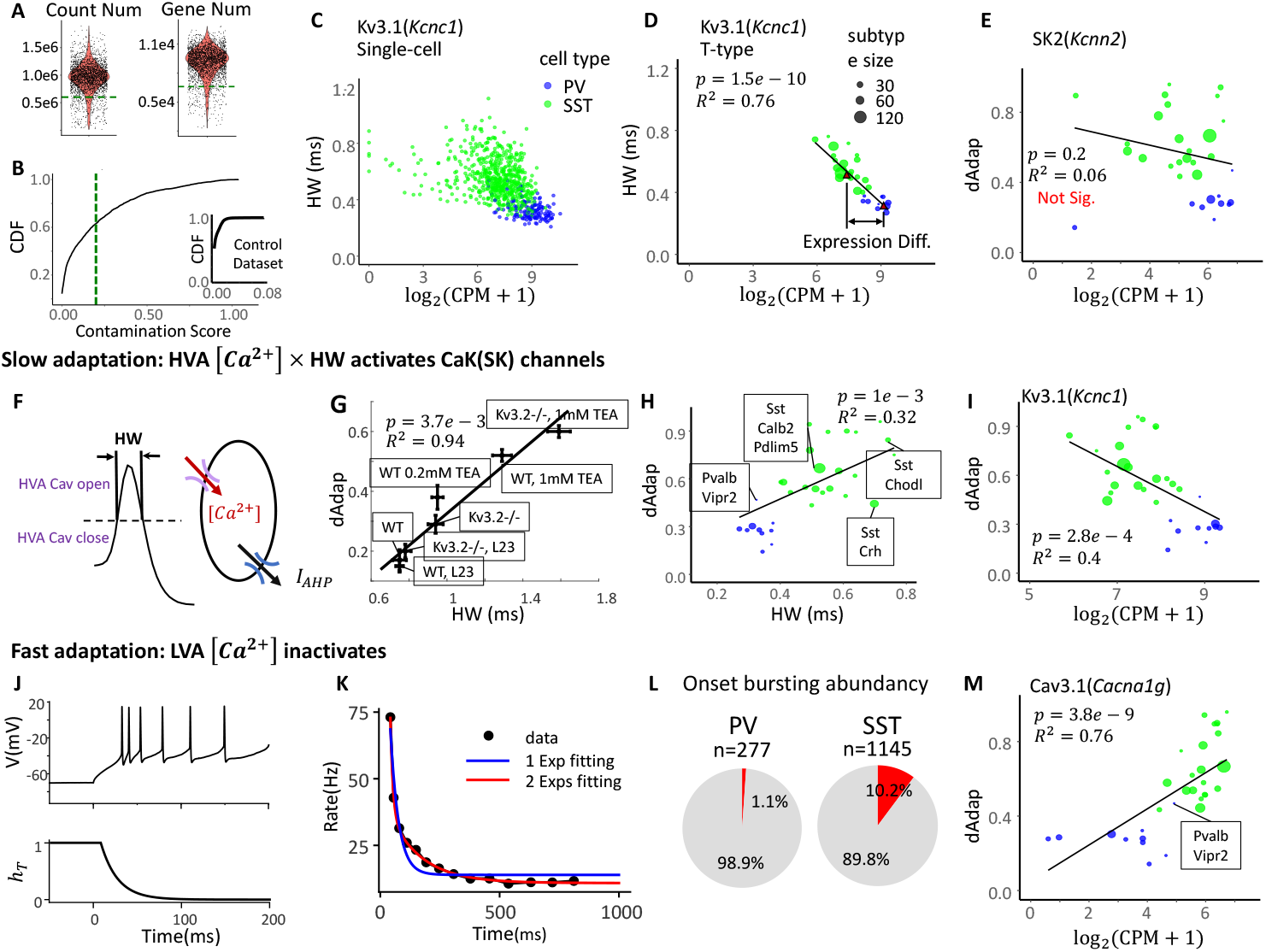
HW and dAdap differences are reflected in the transcriptomic data. (A) Quality control of Patch-seq V1 scRNA-seq data. Dashed lines indicate the filtering thresholds, set just above the tail of the distribution: 6 × 10^5^ (left) and 6,600 (right). (B) Cumulative distribution function (CDF) of the contamination score (CS; see Methods). Cells with CS ¿ 0.2 (dashed line) were excluded. Inset: CS distribution from a dissociation-based dataset^44^, where all cells had CS ¡ 0.08. (C) Scatter plot of spike half-width (HW) versus expression of *Kcnc1* (encoding Kv3.1) across individual V1 PV and SST cells. (D) Bubble plot showing T-type averaged values from (C). The cell-type expression difference is shown by arrows. Dot size indicates sample size; the solid line denotes the weighted linear regression (WLR) fit. *p*: p-value; *R*^2^: proportion of variance explained. (E) Bubble plot of dAdap versus expression of *Kcnn2*. The correlation is not significant. (F) Schematic of the HVA Cav channel mechanism underlying SRA. During each AP, HVA Cav channels open upon sufficient depolarization, with the activation window approximately reflected by HW. The resulting *Ca*^2+^ influx activates SK channels, generating an outward AHP current that contributes to SRA. (G) Increased HW is associated with stronger adaptation. Both HW and dAdap increase following Kv3.2 gene knockout or Kv3 channel inhibition using tetraethylammonium (TEA). Error bars indicate SEM. Data reanalyzed from^31^, collected from layer 5/6 mouse M1 fast-spiking interneurons of either sex at room temperature, unless otherwise noted. (H) Bubble plot of T-type–averaged HW and dAdap in V1. T-type annotation follows^25^. (I) Bubble plot of dAdap versus expression of *Kcnc1*. (J) Voltage depolarization from the current step rapidly inactivates LVA Cav channels, reducing total inward current. (Top) Zoomed-in voltage trace from an example SST cell. (Bottom) Inactivation of the gating variable *h*_*T*_ of LVA Cav channels. (K) Instantaneous firing (IF) rate of an SST cell exhibiting onset-bursting (OB). The IF curve is fitted with either one (blue) or two (red) exponentially decaying components. The two-component fit provides a better match in this case. (L) OB is more prevalent in SST cells (right, 10.2%) than in PV cells (left, 1.1%). (M) Bubble plot of dAdap versus expression of *Cacna1g*, the gene encoding Cav3.1.

Previous work has shown that HW is controlled by voltage-gated K^+^ (Kv3) and Na^+^ (Nav) channels in INs^28,29^. Specifically, higher Kv3 conductance leads to faster repolarization after each spike, resulting in a narrower AP waveform and smaller HW. At the single-cell level, greater expression of *Kcnc1* (encoding Kv3.1) is correlated with smaller HW (Figure 2C), though with substantial variability. Some cells may show zero *Kcnc1* expression due to dropout, where no transcript is detected despite low expression. To enhance signal and mitigate dropout effects, we aggregated cells at the transcriptomically defined type (T-type) level. These T-types, defined via unsupervised clustering across all genes, exhibit consistent electrophysiological and morphological features^25^. Weighted linear regression (WLR) at the T-type level reveals a strong correlation between *Kcnc1* expression and HW (Figure 2D, *p* = 1.5 × 10^−10^). These findings are consistent with prior studies^28,29^ and support the validity of our approach in linking electrophysiological features with transcriptomic data.

Having validated our methodology with HW-related genes, we next examined genes related to dAdap. SRA is commonly attributed to the medium-timescale afterhyperpolarization current (mAHP), mediated by small-conductance Ca-activated K^+^ channels (SK)^22^. These SK channels, activated by Ca^2+^ accumulation during APs, slow neuronal firing and contribute to SRA. We therefore hypothesized that, as with HW, expression of the *Kcnn* gene family would correlate with dAdap. Surprisingly, we found no significant correlation between dAdap and *Kcnn2*, which encodes an SK channel (Figure 2E).

To resolve this discrepancy, we propose alternative mechanisms that may account for the observed differences in dAdap. One contributing pathway involves upstream Ca^2+^ influx through HVA Cav channels, which activates SK channels (Figure 2F). That is, even if SK conductance is similar between PV and SST cells, its activation may differ depending on the amount of Ca^2+^ influx during each AP (Figure 2F). This influx is determined by both the conductance of Cav channels and the duration of their activation. The latter can be approximated by HW, while the former should be reflected in the expression of the corresponding Cav channel genes.

To test this hypothesis, we reanalyze data from^30,31^, which show that manipulating HW, via Kv3.2 knockout or tetraethylammonium (TEA) treatment (an inhibitor of the Kv3 family), modu-lates SRA (Figure 2G). Consistent with this, HW and dAdap are significantly correlated across SST and PV T-types (Figure 2H), and *Kcnc1* (encodes Kv3.1) is also significantly correlated with dAdap (Figure 2I).

In addition, inactivation of LVA Cav channels may also contribute to dAdap on a faster timescale (Figure 2J). Indeed, instantaneous firing (IF) rate curves from some SST cells are better fit by two exponential components rather than a single one, while the latter is the standard for quantifying dAdap (Figure 2K). This fast timescale component can be captured by the initial slope of the IF curve (Supp. Figure 3B). We further classified a cell as having an onset bursting (OB) feature if the fast component is statistically significant and its rectified amplitude exceeds 10Hz (see Methods). Approximately 10% of SST cells exhibit OB, whereas only a few PV cells (1.1%) do (Figure 2L). These OB-positive SST cells are enriched in specific T-types (Supp. Figure 3E). Consistent with this, SST T-types express more *Cacna1g* (encoding LVA T-type Ca channel) than PV T-types (Figure 2M, Supp. Figure 2F), aligning with previous findings^32^.

**Figure 3.**
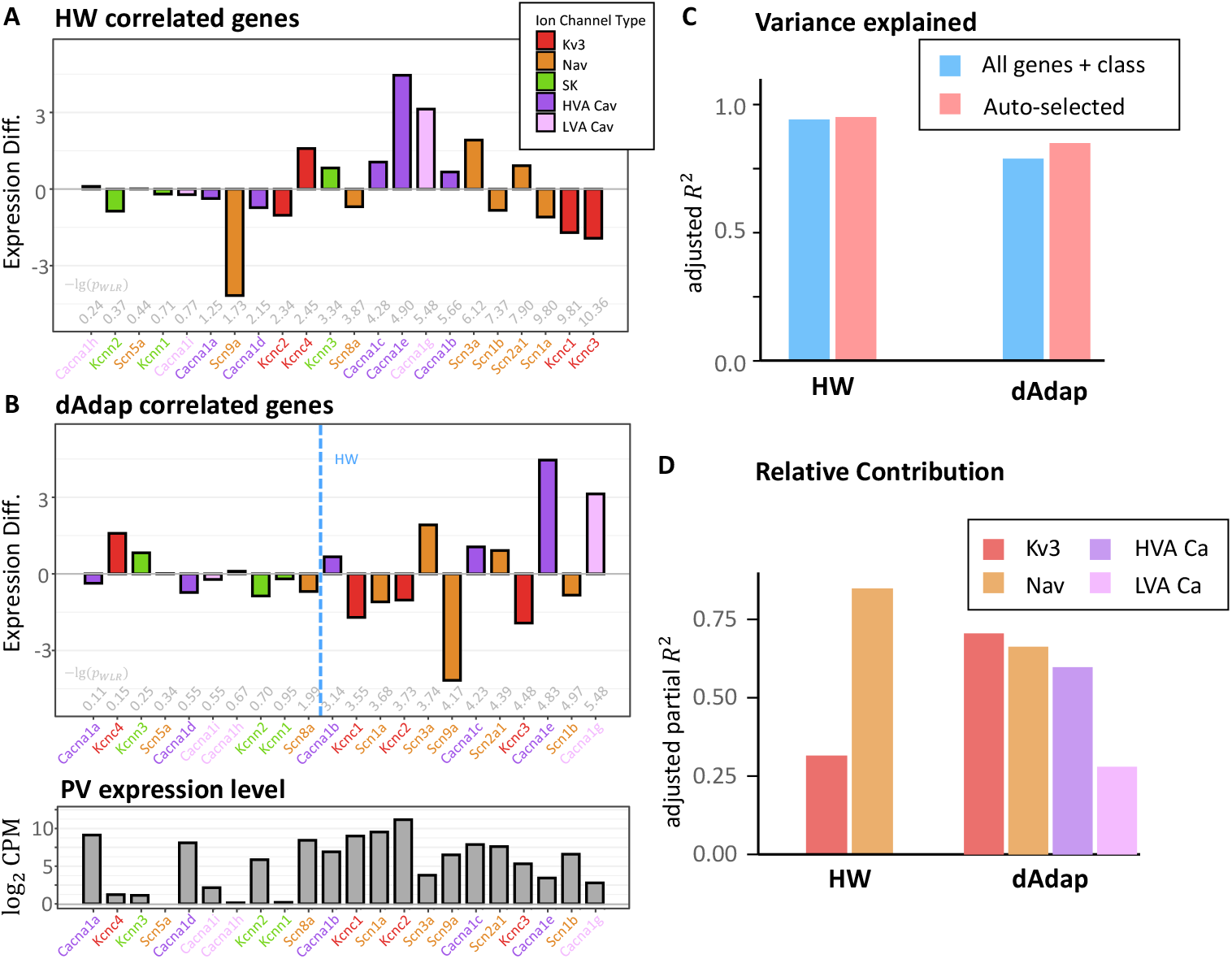
dAdap differences are explained by a combination of gene families. (A) Ranked genes correlated with HW. Genes to the right show stronger correlations with HW, quantified by *p*_WLR_. Significance values (−log_10_(*p*_WLR_)) are annotated in gray. Bar heights represent the expression difference, measured by log_2_ fold change in SST cells relative to PV cells. Color indicates ion channel family. The black dashed line marks *p*_WLR_ < 0.01. (B) (Top) Ranked genes correlated with dAdap. The blue dashed line indicates the correlation significance for HW. (Bottom) Mean gene expression in PV cells. (C) Bar plot of adjusted *R*^2^ from multivariate regression models predicting HW or dAdap from candidate genes. Blue bars: models including all related genes and a class-dependent variable (HW: 94.1%, dAdap: 80.0%; see Methods). Pink bars: models based on auto-selected genes (HW: 95.1%, dAdap: 85.0%). (D) Relative contribution of each ion channel family, quantified by adjusted partial *R*^2^ (see Methods). See Supp. Sheet 2 for further details.

However, our method is limited to qualitatively identifying onset bursting (OB) based on the current data, rather than providing robust quantitative estimates. Typically, only about 3 APs contribute meaningfully to fitting the fast component. Consequently, while the presence or absence of an OB feature can be assessed statistically (see Methods), the amplitude of the fast component exhibits high variability (Supp. Figure 3C, D).

Among the T-types, *Pvalb Vipr2*, which corresponds to chandelier cells, shows elevated dAdap (Figure 2H), consistent with findings from^33^. This T-type also displays strong *Cacna1g* expression (Figure 2L) and includes a cell exhibiting the OB feature (Supp. Sheet 1).

### dAdap differences are explained by a combination of gene families

We next investigate gene families relevant to predicting HW and dAdap, along with their relative importance. We begin by ranking genes based on their correlation significance with HW and dAdap, while showing their expression difference between SST and PV cells (for one example, see Figure 2D). For HW (Figure 3A), the Nav and Kv3 families showed the strongest correlation, consistent with previous findings^28,29^. The corresponding differential expression (DE) analysis is presented in Supp. Figure 4A. Additionally, some gene expressions are highly correlated across IN T-types (Supp. Figure 4B), suggesting they may operate as coordinated functional groups^34^. Moreover, correlation significance is linearly related to the variance explained in the data (Supp. Figure 4C).

**Figure 4.**
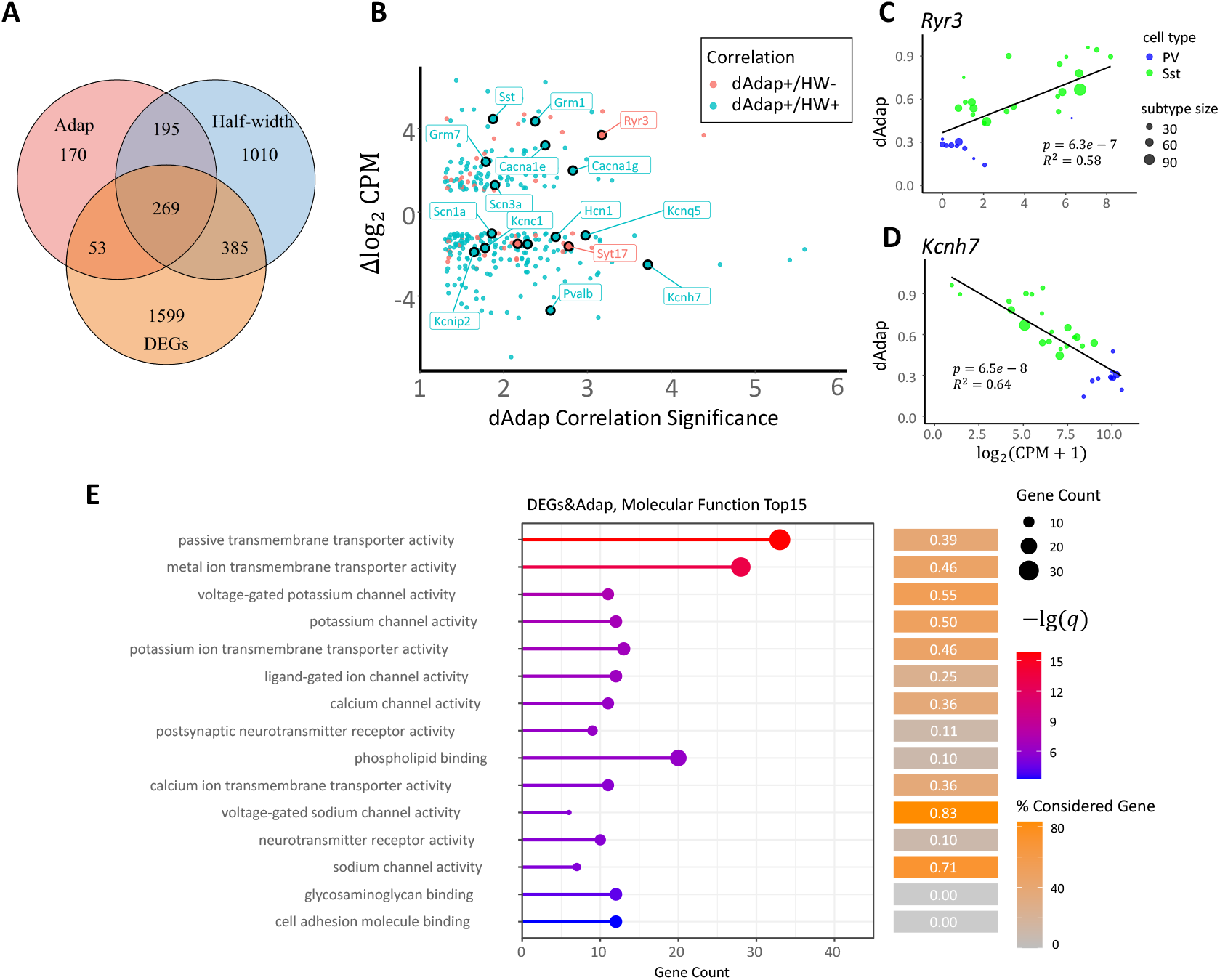
List of potential contributing genes generated by an unsupervised filter algorithm. (A) Venn plot of dAdap highly-correlated genes (pink), HW highly-correlated genes (blue), and differential expression genes (DEGs, orange). (B) Scatter plot of filtered genes. The x-axis indicates the dAdap correlation significance, measured by −*lg*(*q*_WLR_). The y-axis indicates the fold-change compared to PV cells. Genes that are highly correlated with both Adap and HW are labeled in blue, and those that are only highly correlated with Adap are labeled in pink. Here, we study the genes with *q*_WLR_ < 0.05 and *μ*Δ log_2_(CPM)*μ >* 1 (C) Correlation between Adap and *Ryr3*. (D)dAdap and *Kcnh7*. (E) Gene Ontology enrichment analysis of molecular functions. Top 15 terms enriched among genes that are differentially expressed and significantly correlated with dAdap. Colored cubes on the right indicate the proportion of genes that we discussed within each functional group. See Supp. Sheet 3 for details.

Previous work has suggested that SRA is primarily mediated by SK channels^22^. However, none of the SK channel–encoding genes show significant correlation with dAdap differences (Figure 3B, green). In contrast, HW itself (Figure 3B, blue dashed line), HW-related genes (Nav and Kv3 families; orange), and several Cav channel–encoding genes (Figure 3B, purple and pink) are significantly correlated with dAdap. Genes significantly correlated with dAdap tend to be highly expressed in both PV and SST interneurons (Figure 3B, bottom). However, some genes are highly expressed in PV cells yet do not correlate with dAdap differences. For example, *Kcnn2* is highly expressed in PV cells (Figure 3B, bottom), again suggesting the presence of SK channels, but they may not be sufficiently activated by Ca^2+^ influx through HVA Cav channels. Similarly, *Cacna1a* is strongly expressed in both PV and SST cells (Figure 3C). Still, it does not correlate with dAdap, highlighting that a gene may be functionally essential for the mechanism yet not contribute to dAdap variability across cell types.

To evaluate the collective contribution of gene families, we construct multi-gene WLR models including all significantly contributing families (Nav and Kv3 for HW; Nav, Kv3, HVA, and LVA for dAdap), along with a continuous cell-type variable defined by the log_2_ fold change between *Pvalb* and *Sst* expression (see Methods). Together, these genes explain over 90% of the variance (Supp. Sheet 2). However, due to strong inter-gene correlations (Supp. Figure 4B), including all the genes has the risk of overfitting, as the number of predictors is high. To address this, we use adjusted *R*^2^, accounting for model complexity, and implement an auto-selection procedure to retain only the most informative genes (Figure 3C; see Methods). This reduced model improves the adjusted *R*^2^ relative to the full model (Figure 3C). Notably, the continuous cell-type variable is excluded during auto-selection, suggesting minimal cell-type–specific effects in the PV and SST dataset.

Based on the auto-selected model, we quantify the relative importance of each gene family using adjusted partial *R*^2^ (Figure 3D; see Methods). This analysis confirms that all included gene families contribute meaningfully to explaining the observed differences in HW and dAdap. Notably, the LVA Ca^2+^ family (represented here solely by *Cacna1g*) shows the lowest explanatory power, consistent with the observation that only about 10% of SST cells exhibit the OB feature. Interestingly, the LVA Cav gene *Cacna1g* still shows significant correlation with dAdap even after excluding OB cells and OB-enriched T-types (Supp. Figure 5). One possibility is that a small LVA Ca^2+^ current contributes to dAdap but is too weak to generate a significant OB signature detectable by our method.

**Figure 5.**
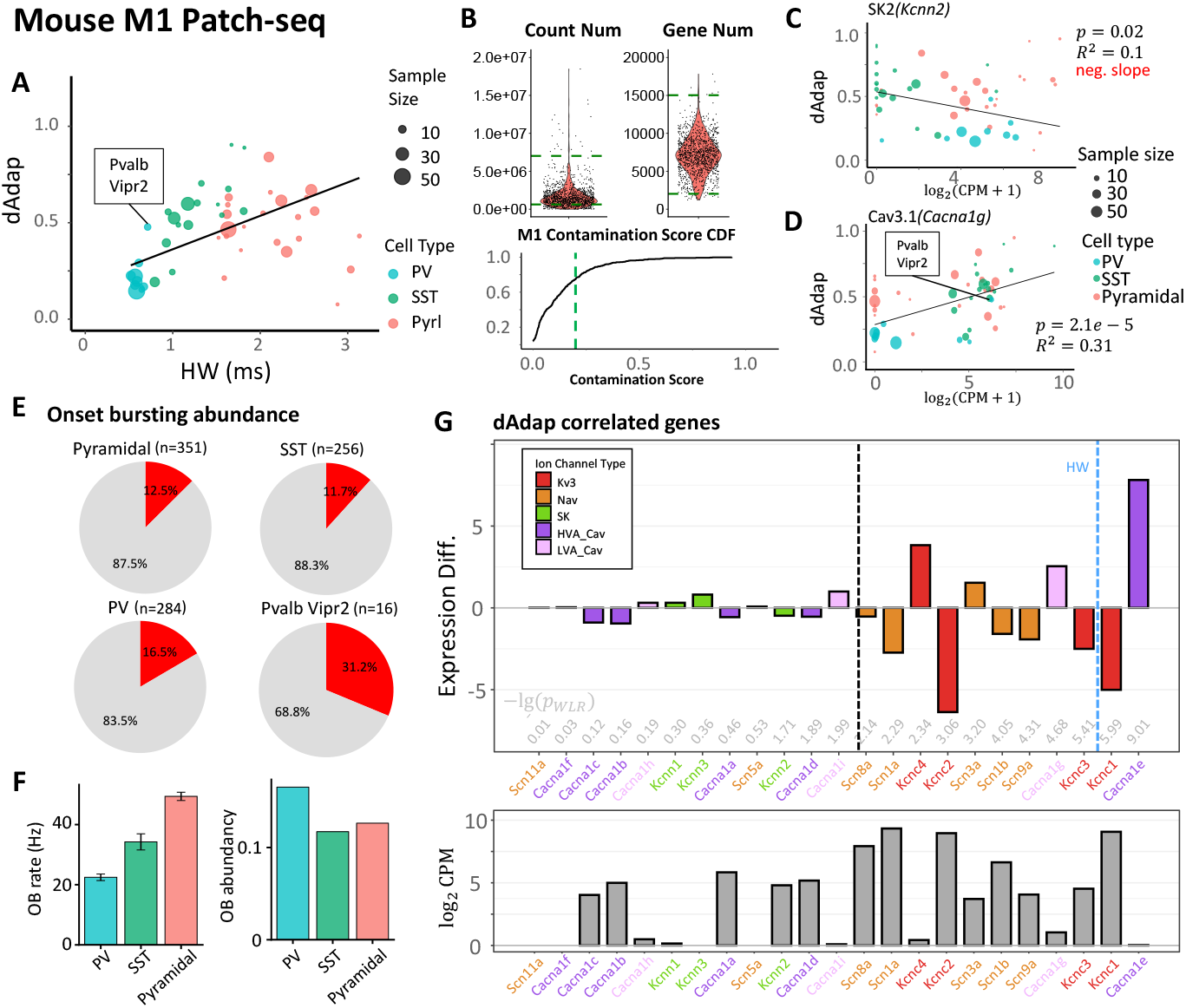
Validation in a mouse M1 dataset. (A) Correlation between HW and dAdap at the T-type level from the mouse M1 dataset. The dashed line indicates the fitting for only PV and SST INs. *Pvalb Vipr2* corresponds to chandelier cells. (B) Quality control for total gene count, gene number, and contamination. (C) Bubble plot for the T-type averaged dAdap and SK2 coding gene *Kcnn2* of M1 Pyramidal, PV, and SST cells. (D) dAdap and Cav3.1 coding gene *Cacna1g*. (E) Pie charts for OB abundance in M1. (F) Bar plot of OB rate (left) and OB abundance (right). However, OB rate measurement is subject to large variance. See Supp.Figure 3. (G) (Top) Ranked genes that correlated with dAdap. The bar heights show the expression difference, measured by the log_2_ fold change of pyramidal cells compared to PV cells. The significance −lg(*p*_WLR_) is annotated in gray. The black dashed line indicates *p*_WLR_ < 0.01. The blue dashed line indicates the correlation significance with HW. (Bottom) Averaged gene expression in PV cells.

Additionally, *Cacna1e*, which encodes the R-type Cav channel (Cav2.3), is significantly correlated with dAdap (Figure 3B; Supp. Figure 4E). However, prior studies have shown that R-type Cav channels may not activate SK channels^35^, suggesting that the observed correlation may reflect an indirect or non-causal correlation.

### Other potential genes

Beyond genes directly linked to our hypothesized mechanisms, other genes may also modulate dAdap. To identify such candidates, we examine differentially expressed genes (DEGs) between PV and SST INs and assess their correlation with dAdap or HW using WLR (Figure 4A; see Methods). In our experience, DEG analysis highlights genes with large fold changes between PV and SST cell types, whereas WLR focuses on whether relative gene expression trends explain observed electrophysiological differences. In total, 322 DEGs are significantly correlated with dAdap; among them, 269 also correlate with HW, while 53 do not.

We further examine the fold change in SST versus PV cells and the false discovery rate (*q*_WLR_) for these 322 genes (Figure 4B). Consistent with our hypothesis, the 269 overlapping genes (blue points in Figure 4B) may influence dAdap by modulating the Ca^2+^ influx window via effects on HW. In contrast, the remaining 53 genes likely affect dAdap by altering Ca^2+^ flow rate or the sensitivity of SK channels to Ca^2+^. However, we cannot fully exclude the possibility that some of these genes are simply PV or SST marker genes, such as *Pvalb* and *Sst*.

Among the resulting gene list (Figure 4B), we identify our hypothesized genes *Kcnc1, Kcnc3, Scn1a*, and *Cacna1g*, supporting the validity of our unsupervised filtering approach. Additionally, *Ryr3*, which encodes the type 3 ryanodine receptor, is significantly more expressed in SST cells (Figure 4C) and is positively correlated with dAdap, but not with HW. Higher *Ryr3* expression has been shown to enhance Ca^2+^ release from intracellular stores^36^, potentially increasing SK channel activation^22^. Other candidate genes include *Grm1*, encoding a group I metabotropic glutamate receptor known to facilitate intracellular Ca^2+^ release, and *Syt17*, which may enable Ca^2+^ binding activity (Supp. Figure 6A).

**Figure 6.**
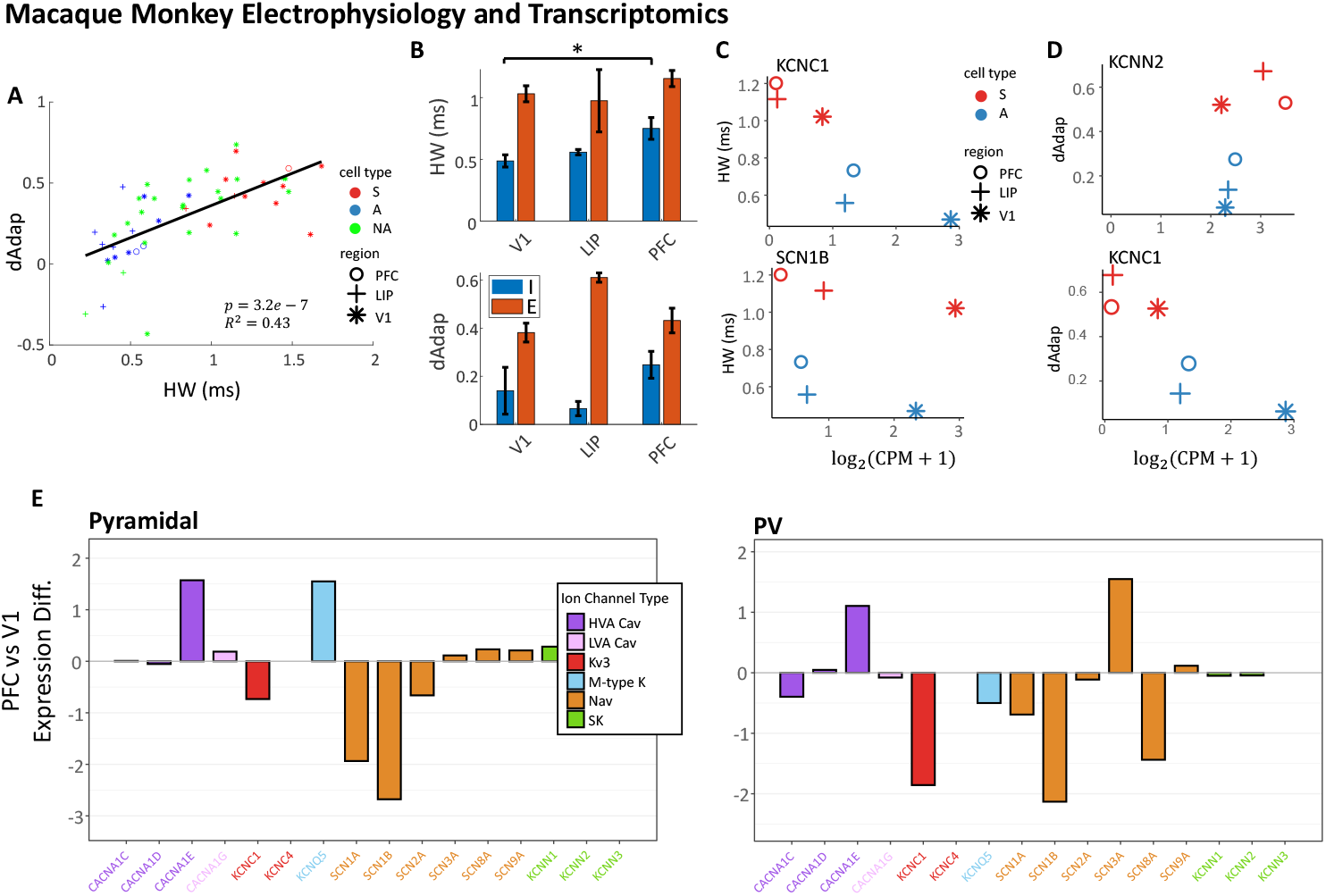
Cell type and regional differences in macaque monkey datasets. Transcriptomic data is from^3^. (A) Correlation between HW and dAdap in a single cell from the multi-regional monkey dataset. Cell types are determined by cell morphology. A: aspiny, putative INs; S: spiny, putative excitatory pyramidal cells; NA: not applicable. Different symbols indicate where the recorded cells are located. *N* = 50. (B) Regional HW (top) and dAdap (bottom) differences in the macaque monkey dataset. *: *p* < 0.05. PFC INs have significantly wider AP and a larger dAdap. *p*_HW_ = 0.02. The error bars indicate standard error. For HW, 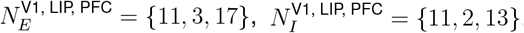.For dAdap, 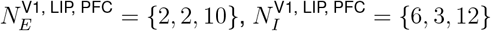 The relationship between HW and expression of *KCNC1* and *SCN1B* across cell types and regions. A: aspiny, supposing INs; S: spiny, supposing excitatory pyramidal cells. Different symbols indicate where the recorded cells are located. (D) same as (C) but for the relationship between dAdap and *KCNN2, CACNA1E*. (E) The expression difference, measured by fold change of gene expressions, in pyramidal cells compared to that in V1 of Pyramidal (top) and PV (bottom) cells. Ion channel families are color-labeled.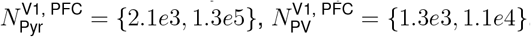. The results from DEG analysis are in Supplementary Sheet 6.

We also identify several K^+^ channel–encoding genes: *Kcnh7, Kcnq5*, and *Kcnip2*, which are significantly more expressed in PV cells and negatively correlated with both HW and dAdap (Figure 4D; Supp. Figure 6B, C). These genes may modulate interneuron excitability^37^. Among them, *Kcnq5* encodes the M-type K^+^ channel, which has been implicated in SRA in CA1 pyramidal neurons^24^. However, if the M-type channel were a primary driver of dAdap, higher *Kcnq5* expression would be expected to associate with greater adaptation. This is contradicted by our data, which show that PV neurons exhibit higher *Kcnq5* expression yet smaller dAdap (Figure 4D).

BK channels may also contribute to the regulation of HW and dAdap (Supp. Figure 6D, E), consistent with previous findings^38,39^. We also observe that the H-current–encoding gene *Hcn1* is significantly differentially expressed between PV and SST cells, although the fold change is close to one. Given that the H-current is activated by hyperpolarization, it is unlikely to influence HW or dAdap under conditions where neurons remain predominantly depolarized.

Finally, the SK channel function is modulated by accessory subunits, including calmodulin, protein kinase CK2, and protein phosphatase 2A^22^. However, the corresponding genes (*Calm1*–*3, Csnk2a1, Csnk2a2, Csnk2b, Ppp2r1a, Ppr2r1b, Ppp2ca, Ppp2cb*) are not DEGs (see Supp. Sheet 3). Thus, these factors are unlikely to explain dAdap differences between cell types. We perform Gene Ontology (GO) enrichment analysis^40,41^ to identify shared functional roles among DEGs, providing biological insight into underlying mechanisms. The GO analysis on our interest 322 DEGs (Figure 4E) reveals enrichment in ion transmembrane transporter activity. Particularly K^+^, Ca^2+^, and Na^+^ channels, which are consistent with our hypothesis. A large proportion of genes identified in the GO analysis have already been discussed (Figure 4E, colored cubes). To the best of our knowledge, the remaining genes among the top 15 GO terms do not show clear functional relevance to SRA. The complete gene list from the GO analysis is provided in Supp. Sheet 4.

### Validation in an M1 Patch-seq dataset

We next ask whether our hypotheses generalize to other brain regions. To test this, we analyze a Patch-seq dataset from the mouse primary motor cortex (M1), in which electrophysiological features were recorded at room temperature^26^. A significant correlation between HW and dAdap is observed (Figure 5A).

After applying similar quality control procedures (Figure 5B) and performing DE analysis (Supp. Figure 7A), we find results qualitatively consistent with the V1 dataset. Specifically, dAdap is not correlated with the SK channel gene *Kcnn2* (Figure 5C), but is correlated with HW-related genes and Cav genes, including *Cacna1g* (Figure 5D). A subset of pyramidal, SST, and PV cells exhibited OB features (Figure 5E). Interestingly, a larger fraction of PV cells displayed OB features compared to the V1 dataset, although the amplitude of the fast component (measured by OB rate) is relatively small (Figure 5F). One potential reason is that neurons in the M1 dataset were more commonly recorded at higher depolarizing currents (e.g., four times the rheobase), making the small-amplitude fast component easier to detect in sweeps with more action potentials. However, differences in recording temperature and brain region may also contribute. The precise mechanism requires further investigation. Some PV T-types (referred to as RNA-types in^26^) were enriched for OB cells. Among them, the *Pvalb Vipr2* T-type, putative chandelier cells, is notably enriched in OB features, consistent with its high *Cacna1g* expression (Figure 5D).

**Figure 7.**
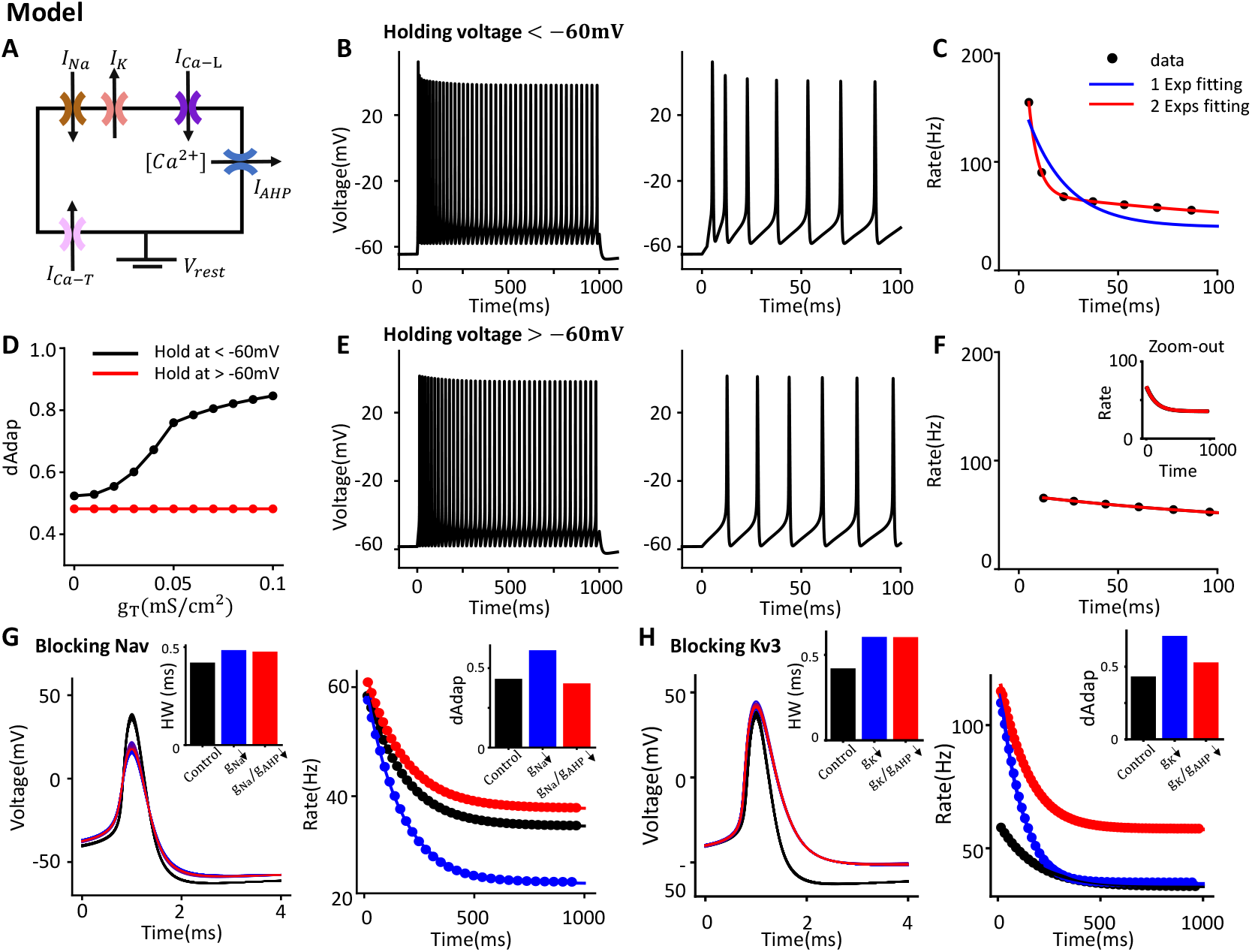
A Hodgkin-Huxley model produces testable predictions. (A) Schematic of the model, which includes five ion channels. Intracellular Ca^2+^ concentration is modeled as proportional to *I*_Ca-L_. (B) Example voltage trace from the model when the holding voltage is below –60mV. (Right) Zoom-in of the first 100ms. (C) Instantaneous firing (IF) curve fit with one versus two exponential components. The two-component fit provides a better match. (D) Model-derived dAdap values under different LVA Ca channel conductance (*g*_*T*_), with holding voltage below –60mV (black) or above –60mV (red). At high holding voltages, the T-type channel is inactivated before the depolarization current. (E, F) Same as (B, C), but with holding voltage above –60mV. In (F), one- and two-component fits are identical, indicating the absence of fast SRA. The trace from 1 to 1000ms is shown as an inset. (G) Predicted changes in HW and dAdap following Nav channel blockade. Three conditions are shown: control ([*g*_*K*_, *g*_*Na*_, *g*_*AHP*_] = [15, 35, 3] mS/cm^2^), reduced *g*_*Na*_ ([15, 25, 3] mS/cm^2^), and reduced *g*_*Na*_ and *g*_*AHP*_ ([15, 25, 2] mS/cm^2^). Insets show HW = [0.42, 0.48, 0.48] and dAdap = [0.43, 0.61, 0.40] for each condition, respectively. (H) Predicted HW and dAdap change by blocking the Kv3 channel. AP waveform and firing-rate adaptation in the control case ([*g*_*K*_, *g*_*Na*_, *g*_*AHP*_] = [15, 35, 3]mS/cm^2^), reduced *g*_*K*_ ([5, 35, 3]mS/cm^2^), and reduced both *g*_*K*_ and *g*_*AHP*_ ([5, 35, 2]mS/cm^2^). The insets show the HW and dAdap in these three cases. The HW of the three conditions are [0.42, 0.61, 0.60], respectively; the dAdap are [0.43, 0.71, 0.53], respectively.

We further analyze which genes are significantly correlated with HW or dAdap (Supp. Figure 7B; Figure 5G). HW correlates with known genes such as *Kcnc1, Kcnc2*, and *Scn1a* (Supp. Figure 7B to D), while dAdap correlates with the same HW-related genes and the Cav gene *Cacna1g*, but not with SK channel genes *Kcnn1*–*Kcnn3*. In addition, *Cacna1e* (Cav2.3, R-type) is significantly correlated with both HW and dAdap, consistent with the previous observation in V1.

However, when analyzing HW and dAdap differences within interneurons and pyramidal cells separately, distinct mechanisms may emerge. This phenomenon is known as the class-driven effect^7^. For instance, across the combined population of excitatory (E) and inhibitory (I) cells, *Kcnc1* and *Scn1a* show a negative correlation with HW (Supp. Figure 7C, D), consistent with the idea that increased K^+^ or Na^+^ current accelerates repolarization or depolarization, leading to narrower AP waveforms. However, when restricting the analysis to either E or I cells alone, this negative correlation persists only within the I population (Supp. Figure 7E, F), highlighting potential class-specific regulatory mechanisms.

We quantify the class-driven effect using the significance of an interaction term in a statistical model, following^7^ (see Methods). This effect is significant for *Kcnc1* in explaining HW differences (Supp. Figure 7E; *p*_class_ = 1.5 × 10^−8^), and is also observed for *Scn1a* (Supp. Figure 7F). The full list of genes with class-driven effects is provided in Supp. Sheet 5. Similar effects are observed for Cav and SK channel–encoding genes (Supp. Figure 7G, H).

We further investigate the mechanisms underlying HW differences in pyramidal cells. None of the Kv3 genes significantly explain the observed HW variability within this population (Supp. Figure 8A). Alternatively, BK channels have been proposed to regulate HW in pyramidal cells^38,39^. However, our data do not reveal a significant negative correlation between BK channel–encoding genes and HW (Supp. Figure 8B, C). Given the low expression levels and pronounced dropout in these genes (Supp. Figure 8A, C), we suspect that the true signal may be obscured by noise, limiting detectability.

### Regional transcriptomic and electrophysiological differences across macaq monkey brain regions

To test whether our hypothesis generalizes across cell types, brain regions, and species, we analyze one electrophysiological and one transcriptomic dataset from macaque monkeys^3^, covering the same three brain areas: V1, lateral intraparietal area (LIP), and prefrontal cortex (PFC). HW and dAdap are measured as before (Supp. Figure 9A, B), with a few cells exhibiting OB features (Supp. Figure 9C).

We first observe a significant correlation between HW and dAdap across individual cells from all three brain areas (Figure 6A). Within each region, both HW and dAdap are consistently lower in interneurons compared to excitatory cells. Across regions, HW increases in pyramidal cells from V1 to PFC, and is significantly higher in PFC than in V1 for interneurons (Figure 6B). Similarly, dAdap is higher in PFC than in V1 for both pyramidal cells and interneurons, though the difference is not statistically significant (Figure 6B).

We next asked whether the observed electrophysiological differences could be explained by transcriptomic data from^3^, which includes whole-brain macaque samples sequenced using unique molecular identifiers (UMIs) and clustered into cell types. For each subtype in each brain region, we calculated a trimmed mean expression. Similar gene expression patterns are observed across macaque PFC, LIP, and V1, consistent with the trends seen in mouse V1. For instance, HW differences are broadly reflected in the expression of *KCNC1* and *SCN1B* (Figure 6C). As in mice, dAdap differences are not explained by *KCNN2* but are instead associated with the HW-related gene *KCNC1* (Figure 6D). These findings suggest that the molecular mechanisms underlying HW and dAdap differences may be conserved between mice and macaques. The full DEG analysis is provided in Supp. Sheet 7.

When comparing across regions (Figure 6E), we find that in pyramidal cells, *KCNC1, SCN1A*, and *SCN1B* all show significant down-regulation in PFC compared to other areas, consistent with the wider HW observed electrophysiologically. Notably, *SCN1A* and *SCN1B* exhibit much larger fold changes than *KCNC1*, suggesting that regional HW differences in pyramidal cells may primarily arise from changes in Na^+^ channels rather than K^+^ channels. Interestingly, both M-type and SK channel–encoding genes show an increasing trend in PFC. Combined with the broader HW, our hypothesis predicts stronger SRA in this region, which aligns with the trend in Figure 6B. We also note increased expression of the M-type channel gene *KCNQ5* in PFC, suggesting a potential role in regional dAdap differences.

INs exhibit a significant increase in HW in PFC compared to V1. This difference could arise from either changes in cell-type composition (e.g., fewer PV cells and more SST cells) or intrinsic property changes across regions. The compositional explanation appears less likely, as PV cells consistently comprise 40–50% of the combined PV and SST population across regions^3^ (though see^21^ for contrasting findings). Nonetheless, single-cell properties may vary unevenly across types. In our analysis, *KCNC1, SCN1A*, and *SCN1B* (but see *SCN3A*) all show significant downregulation in PV INs in PFC (Figure 6E, bottom), consistent with the broader HW observed. Additionally, genes encoding L-type HVA Cav channels and LVA Cav channels show slight downregulation in PFC, while SK channel genes are not differentially expressed across regions. This may help explain why dAdap is not significantly different between PFC and V1. However, due to the lack of statistical significance, this interpretation remains tentative and would benefit from further validation, ideally using a Patch-seq dataset from macaque PFC.

Fold-change comparisons between cell types in V1 and PFC are shown in Supp. Figure 9D.

### A Hodgkin-Huxley model produces testable predictions

To further illustrate the mechanisms of SRA and generate testable predictions, we construct a Hodgkin–Huxley model that incorporates LVA (T-type) and HVA (L-type) Cav channels, as well as SK channels (Figure 7A). The model also includes intracellular calcium dynamics, where the increase in [Ca^2+^] is proportional to the Ca^2+^ current through HVA Cav channels (*I*_Ca-L_), but not LVA Ca channels, consistent with previous findings^23,35^ (see Methods).

Our model first predicts that the presence of fast timescale SRA depends on the holding voltage. We successfully reproduce SRA with both fast and slow components (Figure 7B, C), as demonstrated by fitting the instantaneous firing (IF) curve with one or two exponential decay components (Figure 7D), recapitulating the behavior observed in adaptive SST cells (Figure 2K). Increasing the LVA Ca channel conductance (*g*_*T*_) enhances dAdap (Figure 7E, black line).

Because fast SRA depends on activation of LVA Ca channels, holding the membrane potential above –60mV prior to depolarization should eliminate this component. Consistent with that, depolarizing the holding voltage removed the OB feature in VIP INs^42^, and resulted in a smaller afterdepolarization or delayed depolarization in human neocortical neurons^43^.In our model, when the model neuron is held just above –60mV, T-type channels become fully inactivated before depolarization, resulting in a reduced onset firing rate (Figure 7E, F). Fitting the IF curve with one or two exponential components yields identical results (Figure 7H), indicating the absence of fast timescale SRA. In this condition, dAdap becomes independent of LVA Ca channel conductance (*g*_*T*_) (Figure 7D, red).

Second, we predict that sequentially blocking Nav channels with tetrodotoxin (TTX) and SK channels with apamin can help dissect the mechanisms underlying slow timescale adaptation. To test this, we first eliminate the fast component by setting the LVA Ca channel conductance (*g*_*T*_) to zero. We then reduce the Nav conductance (*g*_*Na*_) and sequentially reduce the SK conductance (*g*_*AHP*_) (Figure 7G). Reducing *g*_*Na*_ reduces the depolarizing current, broadening the AP waveform, and increasing HW. The wider AP allows greater Ca^2+^ influx during each spike, resulting in increased dAdap (Figure 7G, blue). Further reducing *g*_*AHP*_, mimicking the effect of apamin, selectively decreases the SK current without altering the AP waveform. As a result, dAdap decreases while HW remains unchanged (Figure 7G, red).

Our model also predicts that blocking Kv3 channels with tetraethylammonium (TEA) produces a similar effect (Figure 7H).

Finally, we predict that blocking upstream L-type Ca^2+^ channels with toxins such as nimodipine affects dAdap but not HW. To test this, we systematically vary the conductance of Kv3 (*g*_*K*_) and HVA Ca^2+^ channels (*g*_*Ca*−*L*_) (Supp. Figure 10). HW is primarily determined by *g*_*K*_ and remains unaffected by changes in *g*_*Ca*−*L*_ (Supp. Figure 10A), whereas dAdap increases either when *g*_*K*_ is reduced or *g*_*Ca*−*L*_ is increased (Supp. Figure 10B). This dissociation demonstrates that Ca^2+^ influx through L-type channels acts upstream of SK channels to regulate SRA.

## Discussion

Our analysis links transcriptomic differences to AP shape and SRA across subtypes, regions, and species. In mouse V1, HW differences correlate with Kv3 and Nav gene expression, consistent with prior studies. dAdap differences are not explained by SK channels, but rather by upstream Ca^2+^ influx, which is shaped by HW and HVA Cav channel–related genes. Additionally, dAdap is influenced by the onset bursting (OB) feature, which is modulated by LVA Cav conductance. Using an unsupervised filtering approach, we identify additional contributing genes. Similar relationships are observed in mouse M1 and macaque datasets. Finally, we build a minimal Hodgkin–Huxley model with Ca dynamics to generate testable pharmacological predictions.

Our work pioneers the use of new transcriptomic datasets to understand important electrophysiological features of neurons across regions and species. Our methodology becomes available only because of improvements in sequencing technologies. So far, the rich transcriptomic datasets are mostly used in distinguishing different cell types and developing downstream gene-editing tools^3,25,44,45^. However, in many scenarios, this line of study is difficult to unravel the intricate functions of individual genes in explaining neural electrophysiological activities. Early work from^46^ identifies potential genes by combining transcriptomic datasets with electrophysiological datasets. Patch-seq provides electrophysiological and transcriptomic data from the same cell, providing a unique link at an unprecedented level^5^. Only by utilizing this data in the Patchseq can we infer the causal links between gene expression differences and HW and dAdap differences, which can be further tested in other datasets across brain areas and species. We study the SRA because of its important role in many functions, including sensory processing from vision^10^, audition^11^, olfaction^12^, and many others^13,14,16–18^. Based on our analysis, we can test whether the adaptation differences of INs across brain areas lead to functional differences between sensory areas and prefrontal areas in a model. Further, we can infer the strength of IN adaptation across the brain areas based on the multi-regional transcriptomic database and test their functional role in a large-scale model, similar to^47,48^. Naturally, our methodology is not limited to the SRA but can be applied to understand other important features across brain areas, such as bursting, sub-threshold oscillation, etc.

When analyzing SRA, the most intuitive hypothesis is that dAdap differences arise from variations in SK channel conductance between PV and SST INs, given that SK channels mediate the mAHP underlying SRA^22^. However, we find no positive correlation between SK channel–encoding genes and dAdap (Figure 2E). If SK channels were considered in isolation, one might mistakenly conclude they do not contribute to SRA in INs. This highlights the importance of interpreting gene function within the full physiological context, rather than drawing conclusions from a single molecular component.

Further analysis reveals that SK channel activation depends on two upstream factors: the time window and the flow rate of Ca^2+^ influx via HVA Cav channels. Since SST cells have broader APs (i.e., larger HW) than PV cells, they allow more Ca^2+^ influx per AP, even with similar Cav conductance levels. Additionally, we find that *Cacna1g*, encoding an LVA T-type Ca channel, contributes independently to SRA via its fast inactivation following depolarization above –60mV^49^.

In our model, we assume that Ca^2+^ influx through LVA T-type Ca channels does not contribute to SK channel activation. However, a study in rat pyramidal cells^35^ suggests that LVA T-type Ca channels may contribute to the mAHP, albeit to a lesser extent than other Ca channel types. Additionally, *Cacna1e*, which encodes R-type Ca channels, is significantly correlated with dAdap, even though the same study reported no role for R-type channels in mAHP generation. It’s worth noting that their experimental protocol was limited to evoking mAHP with a single action potential over a 50ms window, which may not fully capture the dynamics relevant to SRA measured over a 1-second stimulus train. In addition, the spatial distribution of Ca channels may differ between pyramidal cells and INs, potentially affecting their contribution to SRA. Moreover, the study did not isolate the net effect of T-type channel inactivation, leaving its role unresolved. At last, we cannot fully rule out the possibility that *Cacna1e* and *Cacna1g* may be more of a marker gene in distinguishing PV and SST INs. Further research is needed to clarify the distinct contributions of various Ca channel types to SRA under more physiologically relevant conditions.

Our unsupervised gene analysis revealed additional genes involved in intracellular Ca^2+^ modulation. Among them, *Ryr3* (ryanodine receptor 3) is significantly correlated with dAdap but not HW (Figure 4C), consistent with its known role in mediating Ca^2+^ release from intracellular stores^22,23^. A recent modeling study^50^ also implicates *Ryr3* in adaptation changes in neuropathic pain, further supporting its potential contribution to dAdap regulation. Other mechanisms may involve the M-type K^+^ current^24^, which activates rapidly during depolarization and deactivates slowly below threshold. However, *Kcnq5*, a gene encoding this current, is negatively correlated with dAdap (Supp. Figure 6C), suggesting it may not account for local dAdap differences. Nonetheless, it may contribute to regional differences, as pyramidal cells in PFC exhibit both higher *KCNQ5* expression and greater dAdap compared to those in V1 (Figure 6E).

H-type K^+^ currents have also been linked to mAHP^24^, but likely do not impact SRA in our recordings, as cells are depolarized near threshold during the 1-second stimulus. Interestingly, *Hcn1*, which encodes the H-current, is highly expressed in PV cells (Figure 4B), contradicting prior findings that sag currents—primarily mediated by H-currents—are smaller in PV than SST cells in both mouse visual cortex^51^ and human neocortex^52^. This discrepancy may reflect cell-type–specific differences in the kinetics and voltage dependence of *I*_*h*_ in fast-spiking interneurons^53^.

Finally, we emphasize the importance of handling Patch-seq data with care. On the electrophysiology side, datasets often adopt uniform recording protocols to enable large-scale sampling across brain regions. However, the high variability in cellular responses can obscure key features. For instance, we found that dAdap measurements are only robust when recordings reach at least 1.5× rheobase (Figure 1A). Since PV INs have much higher rheobase values than SST INs, a fixed maximum current injection (e.g., 500pA) may be sufficient for SSTs but inadequate for PVs, introducing sampling bias in electrophysiological feature statistics.

On the transcriptomic side, careful noise control is essential for meaningful interpretation. Because our analysis focuses on a small number of genes involved in the hypothesized mechanisms, we cannot rely on the statistical power afforded by large gene sets typically used in classification studies. To mitigate this, we apply stringent thresholds for total gene counts and feature detection, exclude potentially contaminated cells, and analyze data at the transcriptomic subtype (T-type) level. Additionally, differences between and within cell classes may arise from distinct underlying mechanisms^7^. *Kcnc1*, for example, shows a significant class-dependent effect, indicating divergent regulatory logic in excitatory neurons versus interneurons.

In summary, we show that transcriptomic data can account for differences in AP shape and SRA across cell types and brain regions. Our analysis suggests that expression differences in both HVA and LVA Ca channel–encoding genes, along with HW-related genes, rather than SK channel genes, underlie SRA variability. This underscores the importance of avoiding conclusions based on single-gene analyses. Our methodology offers a framework for predicting electrophysiological properties across brain regions using existing transcriptomic data, paving the way for future experimental and theoretical studies.

### Limitations of the study

One limitation of our study is the exclusive focus on MGE-derived PV and SST INs, excluding CGE-derived subtypes such as VIP and NDNF INs, which together account for approximately 30% of all cortical INs^19^. We excluded CGE-derived INs due to their heterogeneous and often complex firing patterns, such as accelerating and irregular activity^54^, which make the dAdap measurement less reliable. Moreover, limiting our analysis to MGE-derived INs helps reduce potential confounding from class-dependent effects associated with distinct developmental lineages.

## Methods

### Datasets

The transcriptomic data and electrophysiological data were accessed via Allen Institute for Brain Science’s Cell Types Database - [Mouse Patch-seq dataset](https://knowledge.brain-map.org/data/1HEYEW7GMUKWIQW37BO/summary) on January 06, 2020^25^. Electrophysiological recordings were made at physiological temperature (34 °C). This dataset contains 4,284 mouse cells from the primary visual cortex(VISp) layer1, layer2/3, layer4, layer5, layer6a and layer6b. Cells tagged with PV and SST are included in our analysis.

The Patch-seq dataset of adult mouse primary motor cortex (MOp) was accessed from^26^. This dataset collected transcriptomic and electrophysiological data from 1,329 cells at room temperature (25 °C) from mouse MOp layer 1, layer 2/3, layer 5, and layer 6. Cells tagged as PV, SST, and pyramidal (ET, IT, CT, and NP) cell types are included in our analysis.

The electrophysiological dataset from mouse S1 Layer 2/3 was collected from PV-Cre-Ai9 or SST-Cre-Ai9 line as in^55^. Tissue acquisition, processing, and used solutions are described in detail by^56^. The dataset consisted of 16 PV cells and 21 SST cells from 20 individuals of either sex within an age range of 18 days to 35 days.

The macaque monkey transcriptomic dataset, collected through unique molecular identifier (UMI) sequencing, is accessed from^3^. Only the data from monkey No. 2 is used for better area annotation. We exclude the data with ambiguous area annotation, i.e., data from slices across two or more areas. Data from all layers are used.

The intracellular recordings of macaque monkey neurons were obtained in the Inoue and Martinez lab. Tissue acquisition, processing, and used solutions are described in detail by^57^. The dataset consisted of 247 recorded neurons from 11 individuals of both sexes, with an age range of 4.39 to 14.6 years.

### Electrophysiological measurements

We use the same measurement as in^54^. We automatically set up a detection threshold to detect an action potential (AP) in any given voltage trace *V* (*t*) by the following: We first set up an upper bound as *V*_*up*_ = min {max(*V* (*t*)), 0}, then a lower bond as *V*_*low*_ =median(*V* (*t*)). Further, we set the detection threshold as *V*_*thr*_ = max {0.8*V*_*up*_ + 0.2*V*_*low*_, −20}. Next, the *i*th AP is detected at *t*_*i*_ if *V* (*t*_*i*_) < *V*_*thr*_ and *V* (*t*_*i*_ + Δ*t*) ≥ *V*_*thr*_, with the resolution in our recordings Δ*t* = 0.05*ms*. Furthermore, we exclude any APs within 0.5*ms* following another AP. The rheobase current is the minimum current that triggers an AP in the experiments.

The AP-related properties are calculated from aggregated APs from all the sweeps with fewer than 40 APs, while excluding the first AP in each sweep. The maximum voltage of each AP is calculated from the 2*ms* time window following the trace past the AP detection threshold. The AP threshold is calculated as the voltage when the voltage deviation is 20*mV/ms*. The AP half-width is calculated as the time of the AP above the midpoint between the peak and the threshold. To improve precision, linear interpolation is used to calculate the threshold and AP half-width.

The degree of adaptation (dAdap) is calculated from the instantaneous firing (IF) curve, excluding the first AP from each recording sweep. IF curves are calculated as *r*(*t*_*i*_) = 1*/*(*t*_*i*+1_ − *t*_*i*_). Next, we fit the IF curve to an exponential function *f* (*x*) = *a* + *b*exp(−*cx*), using *curve fit* function of *scipy* package in Python. The timescale of the fitting is required to be larger than 20ms (equivalently *c* <= 50*Hz*). The dAdap is the proportion of the change in firing rate between 0s and 1s: dAdap = 1 − *f* (1*s*)*/f* (0). Due to the large variance of dAdap around the rheobase, we choose the dAdap from the first sweep above 1.5 times the rheobase. In addition, the qualified sweep must be recorded below the 4 times rheobase, the last spike must occur later than 0.8 seconds, and the sweep must have at least 10 APs. Otherwise, the cell will be excluded from further analysis.

To quantify whether a cell has an onset bursting (OB) feature, we fit the IF curves to a function with two exponential components *f* (*x*) = *a* + *b*_1_exp(−*c*_1_*x*) + *b*_2_exp(−*c*_2_*x*). During the fitting, the timescale of the fast component is required to be less than 20*ms* (or equivalently *c*_1_ > 50*Hz*) and the slow timescale is required to be larger than 20ms (equivalently *c*_2_ <= 50*Hz*). The fast amplitude *b*_1_ is referred to as the rectified onset bursting rate. The significance of OB of a sweep is determined by the p-value of the fast amplitude *b*_1_ by the following equation *p* = 2(1 − *t*_cdf_(|*b*_1_.*μ*|/*b*_1_.*σ*), *df*) where *t*_cdf_ is the cumulative probability of student’s t-distribution. *b*.*µ* and *b*.*σ* are the mean and standard deviation of fitted parameter *b*_1_, *df* is the degree of freedom. The fitting is done using the Python package lmfit.

The significance of a cell is determined by the minimum adjusted p-value of all the recorded sweeps by the number of sweeps through the BH correction. The corresponding OB rate *b*_1_ is considered the OB rate for the cell. A cell is considered to have the OB feature if the p-value is less than 0.05 and the rectified OB rate *b*_1_ is larger than 10*Hz*. The OB abundance is determined by the number of cells showing a significant q-value *q* < 0.05 that is adjusted by the cell number of the corresponding T-type.

We investigate the effect by removing cells with an OB feature. To do that, we either remove the cells that have an OB feature, or remove the T-type with OB abundance larger than 20%. See supplementary Sheet 1 for details.

### Quality control of the transcriptomic data

From the raw counts in the Patch-seq database, we exclude cells whose number of unique sequenced genes is less than 6600 and cells whose total counts are less than 6 ×10^5^ (Figure 2 B, Supplement Figure 7A). Genes sequenced in less than 100 cells were filtered out. Raw counts were normalized by count per million (CPM) and then transformed to log_2_(CPM + 1). We refer to a dissociated single-cell dataset^44^ as the uncontaminated database. Following^27^, the contamination score *CS* describes how much microglial marker genes^58^ are present in the sequenced cells.

Single-cell RNA samples collected by Patch-seq could be contaminated by neighboring microglial cells, but not those collected by dissociated cells

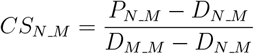

Here, *X*_*A*_*_*_*M*_ represents the median of 50 marker genes from the Patch-seq (P) or dissociated (D)dataset of type *A* in microglial cells. The marker genes differ across cell types (PV or SST IN, pyramidal cell, microglial cell). We adopt the marker gene list for each cell type from^44^. The contamination score *CS* ranges from 0 to 1, reflecting the excess of off-target marker expression. We limited our analysis to cells with *CS* < 0.2 (Figure 2 C, Figure 5 B)

### Classifier

For training a classifier based on HW and dAdap, we find the best linear manifold that splits the cells by using the Support Vector Machine algorithm in Matlab. The resulting line is indicated by *β*_1_*x* + *β*_2_*y* + *b* = 0. To test the robustness of this method, we randomly split the data points into 80% training set and 20% testing set. We calculate the average performance of 100 repetitions < Accy_test_ >. To identify cells with low classification accuracy, we employed a Gaussian Mixture Model (GMM). This model assumes that the data from a cell type are generated from a mixture of two Gaussian distributions, calculated by Matlab’s *fitgmdist* function. The classification of each data point was based on the highest posterior probability derived from the posterior function, which computes the likelihood that each data point belongs to each Gaussian component. The curves with 95% posterior probability for each cluster are reported as confidence boundaries.

The fitted parameters are included in the Supp. Spreadsheet 1.

### Multi-gene linear regression model

Some genes, though passing QC in the published dataset, exhibit zero counts in many cells—a common issue known as dropout. This is illustrated in Supp. Figure 2D, F. A zero count may reflect either truly low expression or a detection failure in a cell that does express the gene. To minimize the impact of this ambiguity, we quantify the expression prevalence for each gene, defined as the number of cells with non-zero counts. We then exclude genes in the bottom 10th percentile of the expression prevalence distribution from subsequent analyses.

Each cell in the dataset is associated with a T-type tag. The mean trimmed expression for each T-type is calculated by averaging the expression of individual cells after removing the top and bottom 25% expression.

We used a weighted linear regression to study the correlation between HW or dAdap with different genes. The significance of observing a non-zero slope *p* and the variance explained *R*^2^ are reported for each pair. When comparing multiple models, correction for the multiple testing problem is necessary. To do so, we adjusted p-values with the false discovery rate (FDR) using the Benjamini-Hochberg method^59^ (Supp. Figure 4B).

We train the multiple linear regression model to predict HW or dAdap based on genes. We first test the performance of the prediction by including all the significant correlated genes identified in Figure 3.

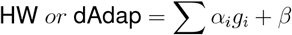

where *g*_*i*_ is the log_2_ of a gene, and *β* is the intercept. The weights *α*_*i*_ are fitted through a multiple variable linear regression. Noticing that we only have 31 T-type (10 PV and 21 SST), the more genes we include, the more likely we are to observe an overfitting problem. We next adapt the Akaike Information Criterion (AIC) to select the model that explains the greatest variance with the fewest number of genes^60,61^. Furthermore, since we hypothesized that dAdap depends on HW, the performance of predicting dAdap using HW and *Cacna1g* is tested.

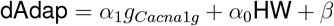

The genes and corresponding fitted weights are listed in the Supp. Spreadsheet 3 file.

After multi-linear regression and auto-selection based on AIC, we next calculated partial R^2^. To quantify the unique contribution of specific gene families within the auto-selected multiregression model, we computed both the partial R^2^ and the adjusted partial R^2^ for each gene family.

Compared to the auto-selected model, we define reduced models by removing the genes that belong to different gene families. The partial R^2^ was computed as the proportion of residual sum of squares (RSS) reduction when moving from the reduced model to the full model:

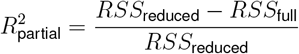

where *RSS*_reduced_ is the residual sum of squares from the reduced model (excluding the term(s) of interest), and *RSS*_full_ is the residual sum of squares from the full model.

This value reflects the additional variance explained by the removed genes, beyond what is already explained by the remained genes.

To account for model complexity and sample size, we additionally computed adjusted partial R^2^ using the following formula:

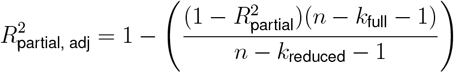

where *n* is the number of observations (sample size), same for both models, and *k*_*full*_ and *k*_*reduced*_ are the number of estimated coefficients excluding the intercept in the full and reduced models, respectively. This adjusted measure penalizes for differences in model complexity, providing a more conservative estimate of unique explanatory power.

### Regression respect to Cell Class

To test which genes have a class-driven effect in explaining HW or dAdap, we fit a class-driven linear model as in^7^:

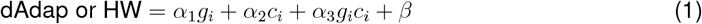

where *g*_*i*_ is the log_2_(*CPM* + 1) of a T-type, *c*_*i*_ is a categorical variable that indicates the class for the *i*th T-type. The significance of the class-driven effect is indicated by the adjusted p-value for *α*_3_, calculated by the ANOVA test. Considering this class-driven effect, we further reported p-values for pyramidal cells and interneurons separately by fitting linear models to the pyramidal cell data and the interneuron data, respectively. The details of the analysis are listed in the Supp. Sheet 5 file.

### Differential expression analysis

Differential expression analysis was performed with R 4.2.3 and the DESeq2 package in R (1.36.0)^62^. In analyzing Patch-seq mouse data, genes with q-value < 0.05 and log_2_(foldchange) > 1 are considered differentially expressed genes (DEG). In analyzing UMI macaque monkey data, genes with q-value < 0.01 and log_2_(foldchange) > 0.4 are considered differentially expressed genes (DEG). In estimating the size factor of the macaque monkey, we used the “poscount” option offered by the DESeq2 package to process the data with lots of zero readouts. Otherwise, we use the default parameters offered by the DESeq2 package.

### Gene Ontology Enrichment

For selected genes, Gene Ontology^40,41^ enrichment analysis is performed using the R package clusterProfiler (4.4.4) with the over-representation analysis (ORA) approach^63,64^.

### Hodgkin-Huxley model with Calcium dynamics

We build a Hodgkin-Huxley neuron model equipped with *Ca*^2+^ dynamics, low-threshold and high-threshold voltage-gated *Ca*^2+^ channels corresponding to T-type and L-type calcium channels, respectively, and *Ca*^2+^ activated *K*^+^ ion channels to reproduce the observed heterogeneity spike-width and degree of adaptation in interneurons. The equations are the following:

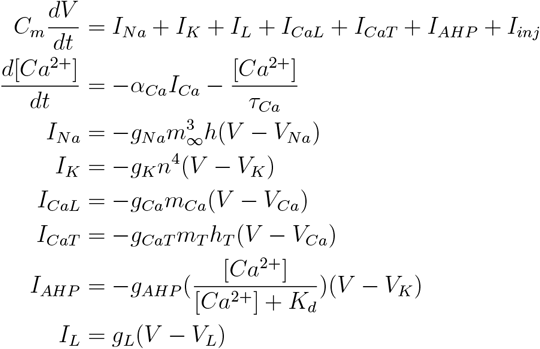

Within the equations, *g*_*Na*_,*g*_*K*_,*g*_*CaT*_, *g*_*CaL*_, *g*_*AHP*_ and *g*_*L*_ are the conductance for the voltage-gated sodium, potassium, low-threshold calcium, high-threshold calcium channels, and the calcium-activated potassium channels, respectively; *V*_*Na*_,*V*_*K*_,*V*_*Ca*_, and *V*_*L*_ are the reversal potentials for sodium, potassium, calcium, and the leak, respectively.

*I*_*Na*_ and *I*_*K*_ are adopted from^65^, with all voltage dependencies shifted by +10 mV to match the experimentally observed reset voltage. This adjustment preserves the original AP dynamics while aligning the model more closely with our data.

The gating variables *m, h* and *n* satisfy the first-order kinetics given by:

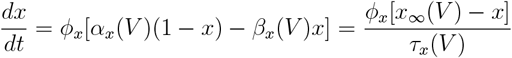

Here, ϕ_*x*_ = 5 is a temperature factor. For 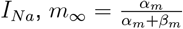 where 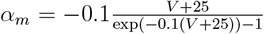 and 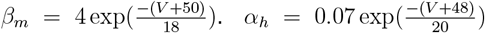 and 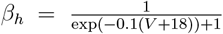.For 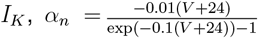 and 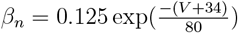.

We adopt the low-threshold *Ca*^2+^ dynamics from^66^, where *m*_*T*,∞_ = *H*(*V* − *V*_*h*_) where *H* is the Heaviside step function and *V*_*h*_ is the threshold responsible for the activation of burst spiking. The inactivation variable *h* has dynamics given by:

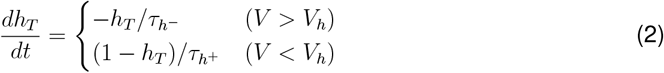

We then adopt the high-threshold *Ca*^2+^ dynamics and adaptation current *I*_*AHP*_ through the *Ca*^2+^ activated SK channel from^67^, where 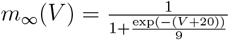

We use the following parameter values unless otherwise specified: *C*_*m*_ = 1*µ*F*/*cm^2^, *g*_*L*_ = 0.1mS*/*cm^2^, *g*_*Na*_ = 35mS*/*cm^2^, *g*_*K*_ = 9mS*/*cm^2^, *g*_*CaL*_ = 0.1mS*/*cm^2^, *g*_*CaT*_ = 0.05mS*/*cm^2^ *g*_*AHP*_ = 3mS*/*cm^2^, *K*_*d*_ = 30*µ*M, τ_*h*−_ = 5ms, τ_*h*+_ = 100ms, τ_*Ca*_ = 200ms, *α*_*Ca*_ = 0.002*µ*M*/*(ms *µ*A), and reversal potentials *V*_*L*_ = − 55mV, *V*_*Na*_ = 65mV, *V*_*K*_ = − 80mV, *V*_*h*_ = − 60mV, *V*_*Ca*_ = 120mV,

The dAdap is always measured from the sweep with a depolarization current injection of *I*_*inj*_ = 2*µ*A*/*cm^2^. The default holding current is 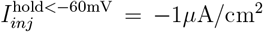. To test conditions under which the T-type Ca channel is inactivated, the holding current is raised to 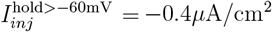.

## RESOURCE AVAILABILITY

### Lead contact

Requests for further information and resources should be directed to and will be fulfilled by the lead contact, Xiao-Jing Wang.

### Materials availability

This study did not generate new materials.

## Supporting information

Sheets S4 Raw data generated from V1 GO analysis. Related to Figure 4.

Sheets S5 Aggregated M1 data at T-type level. Related to Figure 5.

Sheets S6 M1 weighted linear regression result. Related to Figure 5.

Sheets S7 DESeq analysis results on Macaque data. Related to Figure 6.

Sheets S1 Aggregated V1 data at T-type level. Related to Figure 2.

Sheets S2 Parameters for multiple variable linear regression. Related to Figure 3.

Sheets S3 Raw data generated from the unsupervised filtering algorithm. Related to Figure 4.

## Data and code availability

- All data reported in this paper are included in the supplementary sheets.
- All original code has been deposited at GitHub https://github.com/johnhongyumeng/Patchseq and is publicly available as of the date of publication.
- Any additional information required to reanalyze the data reported in this paper is available from the lead contact upon request.

## Acknowledgement

We thank Zhenghan Liao, Panagiota Theodoni, Löic Magrou, and other members of Xiao-Jing Wang lab for the discussion. We thank Shlomo Dellal, Robert Machold, Chiung-Yin Chung, and other members of Bernardo Rudy lab for the discussion. We thank Shreejoy Tripath for his suggestions and feedback on this work. We thank Julia Sunstrum, Sam Mester, Meghan Wiederman, Sara Matovic, Michelle Jimenez, and Michael Poulter for the electrophysiological data collection of the macaque monkeys. We thank Maximiliano José Nigro for sharing mouse S1 L23 data. We thank Yidi Sun and Juan Meng from ION, Shanghai, for sharing the transcriptomic data of the macaque monkey and for their help with the technique. This work is supported by NIH grant R01MH062349, ONR grant N00014-23-1-2040 and the James Simons Foundation grant NC-GB-CULM-00003138 (to XJW), Neuronex NSF 2015276 (to Amy Arnsten and XJW), NIH grants P01NS074972 and R01NS133751 (to BR), CIHR, NSERC, Autism Research Chair province of Ontario, NEURONEX (to JMT), the Canadian Institute for Health Research (NGN 171424) in relationship to Neuronex NSF 2015276 and BrainsCAN accelerator grant funded by a Canada First Research Excellence Fund (to WI).

## AUTHOR CONTRIBUTIONS

Conceptualization, J.H.M., B.R., and X.-J.W.; methodology, J.H.M., Y.K., W.I., J.M.T., B.R., and X.-J.W.; investigation, J.H.M., Y.K., A.L. and M.F.; writing-–original draft, J.H.M and Y.K.; writing-–review & editing, J.H.M, Y.K., B.R. and X.-J.W.; funding acquisition, W.I., J.M.T., B.R., and X.-J.W.; supervision, J.M.T., B.R., and X.-J.W.

## DECLARATION OF GENERATIVE AI AND AI-ASSISTED TECHNOLOGIES

During the preparation of this work, the authors used ChatGPT-4o in order to assist with proofreading and improving the clarity of the manuscript. After using this tool, the authors reviewed and edited the content as needed and take full responsibility for the content of the publication.

## SUPPLEMENTAL INFORMATION INDEX

Figures S1–S10 and their legends in a PDF

Sheets S1 Aggregated V1 data at T-type level. Related to Figure 2.

Sheets S2 Parameters for multiple variable linear regression. Related to Figure 3.

Sheets S3 Raw data generated from the unsupervised filtering algorithm. Related to Figure 4. Sheets S4 Raw data generated from V1 GO analysis. Related to Figure 4.

Sheets S5 Aggregated M1 data at T-type level. Related to Figure 5.

Sheets S6 M1 weighted linear regression result. Related to Figure 5.

Sheets S7 DESeq analysis results on Macaque data. Related to Figure 6.

**Supp. Figure S1.**
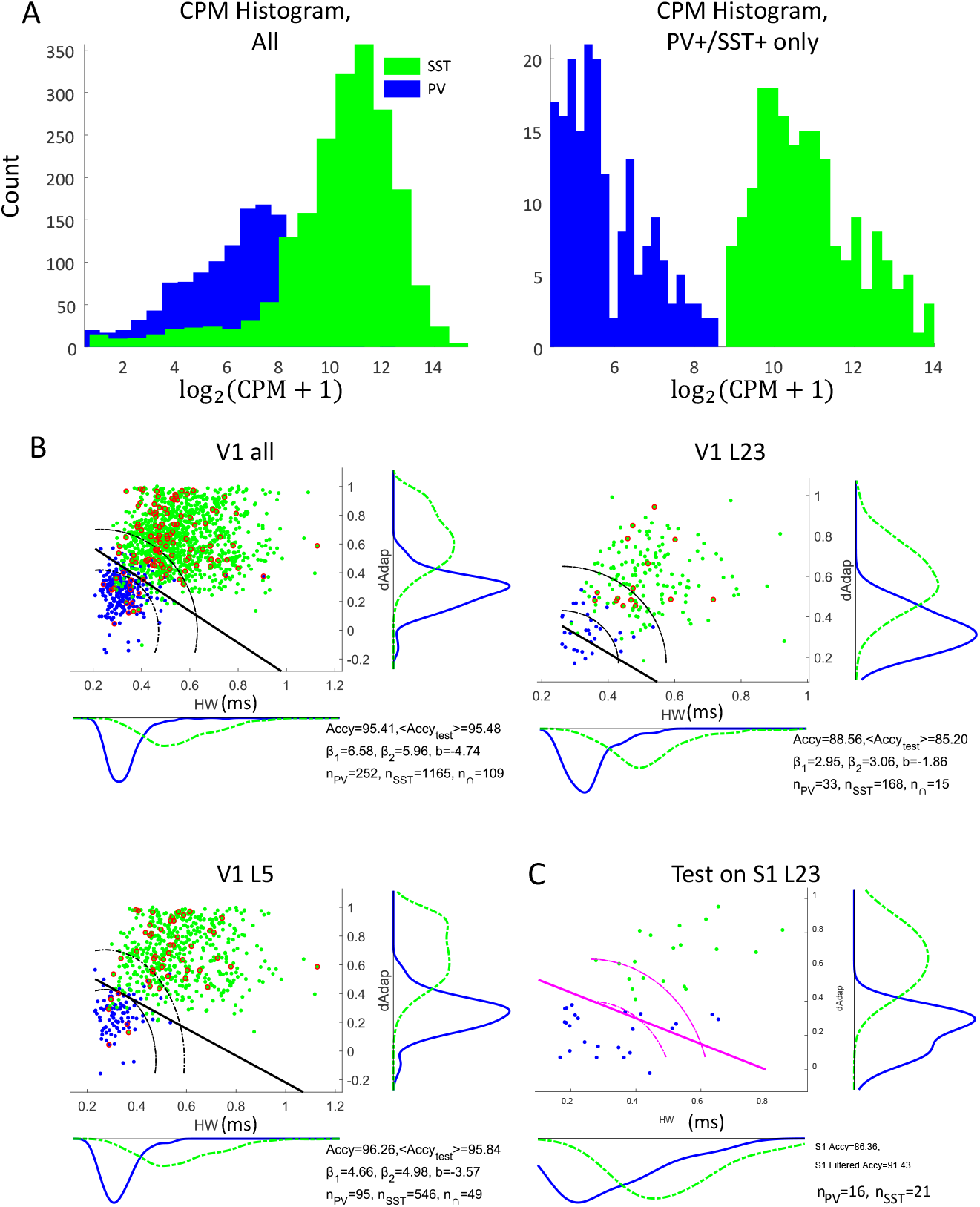
Performance of the trained classifier compared to that of the marker gene classification. Related to Figure 1. (A) distribution of the *Pvalb* and *Sst* markers. Left: Histogram of *Pvalb* and *Sst* in all cells. Right: Histogram of cells that are both *Pvalb*+ and *Sst* +. A cell with *PV* larger than the 20th percentile of the total distribution (*θ*_*PV*_ = 4.49) counts as *Pvalb*+. The same applies for *Sst* + (*θ*_*SST*_ = 8.97). In the dataset, a total of 1025 *Pvalb*+ cells and 1604 *SST*+ cells are identified. Within those, 189 cells are identified as *Pvalb*+/*Sst* +. The CPM distributions of these 189 cells are shown on the right. If considering multiple-gene classification as the ground truth, then the accuracy of the gene-marker (PV and SST marker) classification is 89.62%. (B) Classification of PV and SST based on HW and dAdap across layers. The solid line represents the classification boundary, indicated by *y* =−(*xβ*_1_ + *b*)*/β*_2_. The red circle represents *Pvalb*+/*Sst* + cells. Notice that these cells are not only distributed close to the boundary. Dashed line, the 95% confidence boundary based on the Gaussian mixture model. See Supp. Spreadsheet 1 for details. (C) Test the trained classifier from V1 on a dataset from the somatosensory cortex. The filtered accuracy represents the classifier accuracy after excluding low-confidence cells. I.e., excluding the cells within the two dashed pink lines.

**Supp. Figure S2.**
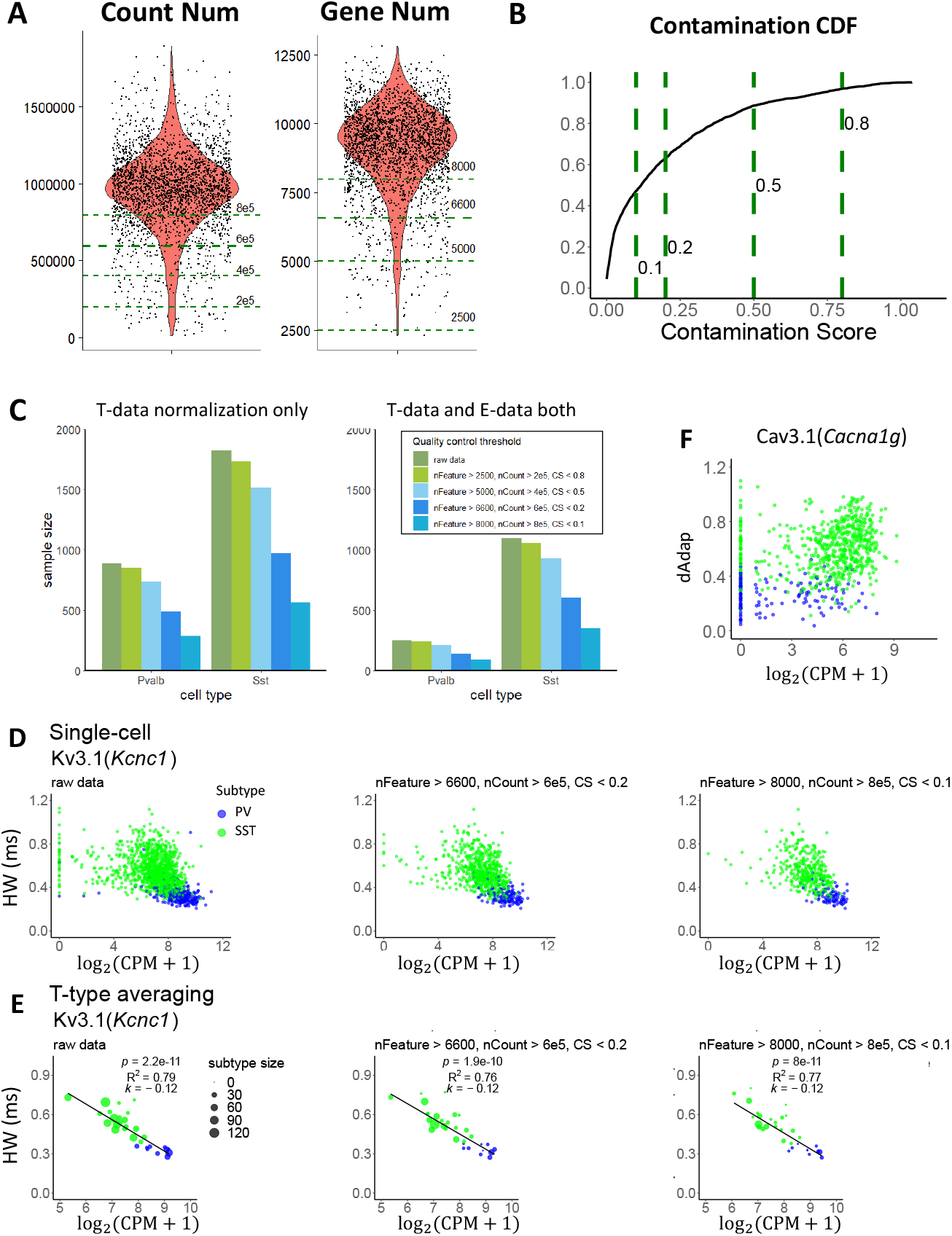
Quality control of transcriptomic data with different thresholds. Related to Figure 2. (A) Violin plot for V1 raw count data: number of total raw counts per cell (nCount RNA, left) and number of unique sequenced genes per cell (nFeature RNA, right). Dashed lines represent different thresholds. (B) Cumulative distribution function for the contamination score. (C) Histogram of the number of cells (sample size) that passed the corresponding quality control parameters before (left) and after (right) merging with cells that also passed the electrophysiological quality control. The parameter sets are colored. (D, E) Scatter plot at the single cell level (D) and at the T-type level (E) of the HW and *Kcnc1* with different quality control parameters. Strict quality control reduces dropout issues (zero readouts along the y-axis) at the single-cell level, while T-type level performances are comparable. (F) Scatter plot at the single cell level of the dAdap and *Cacna1g* after the default quality control. For low-expression genes, the dropout issue remains obvious.

**Supp. Figure S3.**
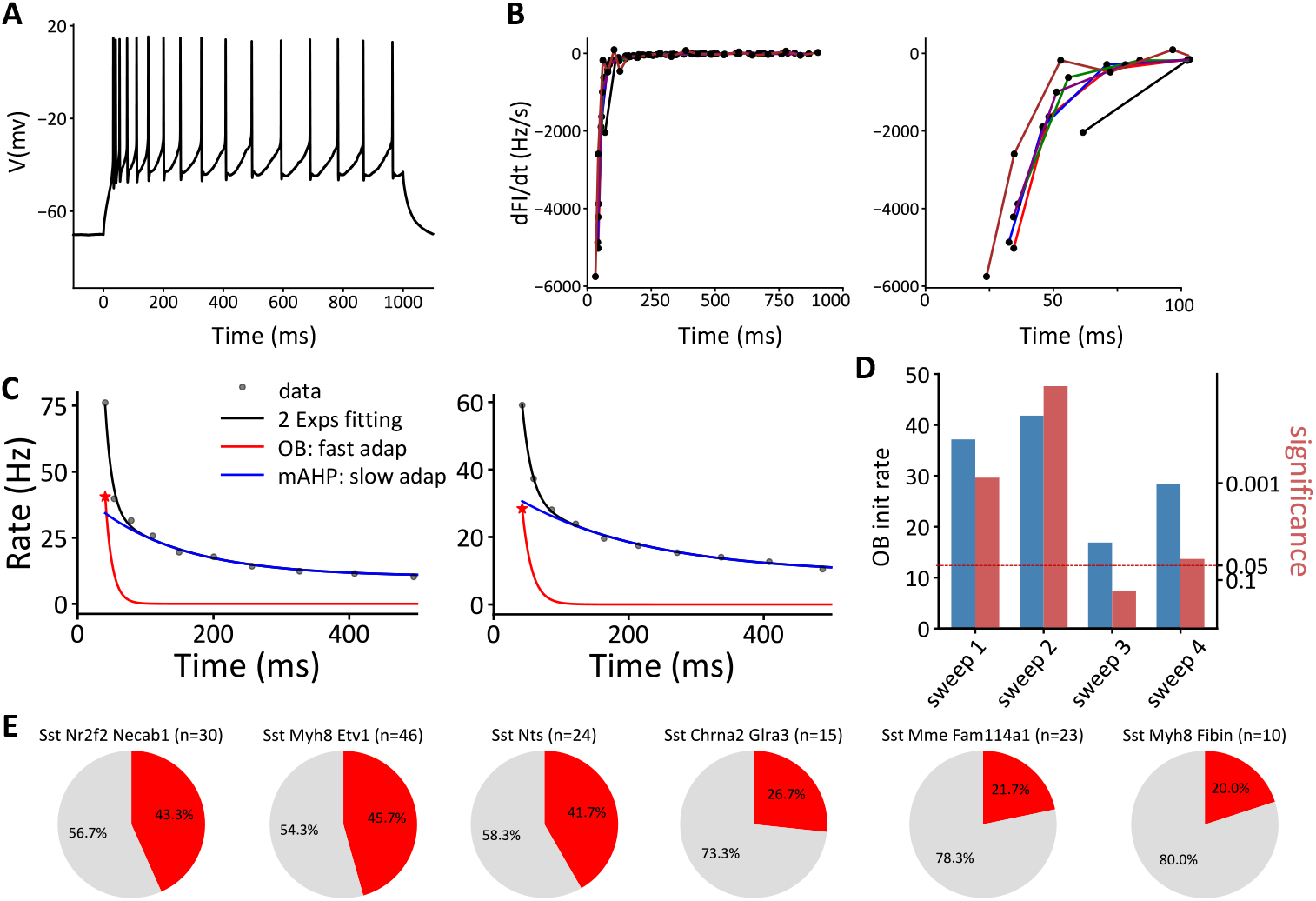
Analysis of the onset bursting feature. Related to Figure 2. (A) Example cell (same as in Figure 2J, K), recorded four times at the same depolarizing current. (B) Instantaneous firing rate (IF) curves across sweeps. Right panel: zoomed-in view. Different colors represent different sweeps. (C) Example IF curve fitted with two exponential components. Color indicates different components of the fitting. Red indicates the fast component of the fitting. (D) Fitted results across the four sweeps. Although three of the four sweeps show a significant OB feature, the initial OB rate varies substantially, making it unsuitable for quantitative analysis. (E) Cells with OB features are enriched in specific T-types. See Supp. Sheet 1 for the full list.

**Supp. Figure S4.**
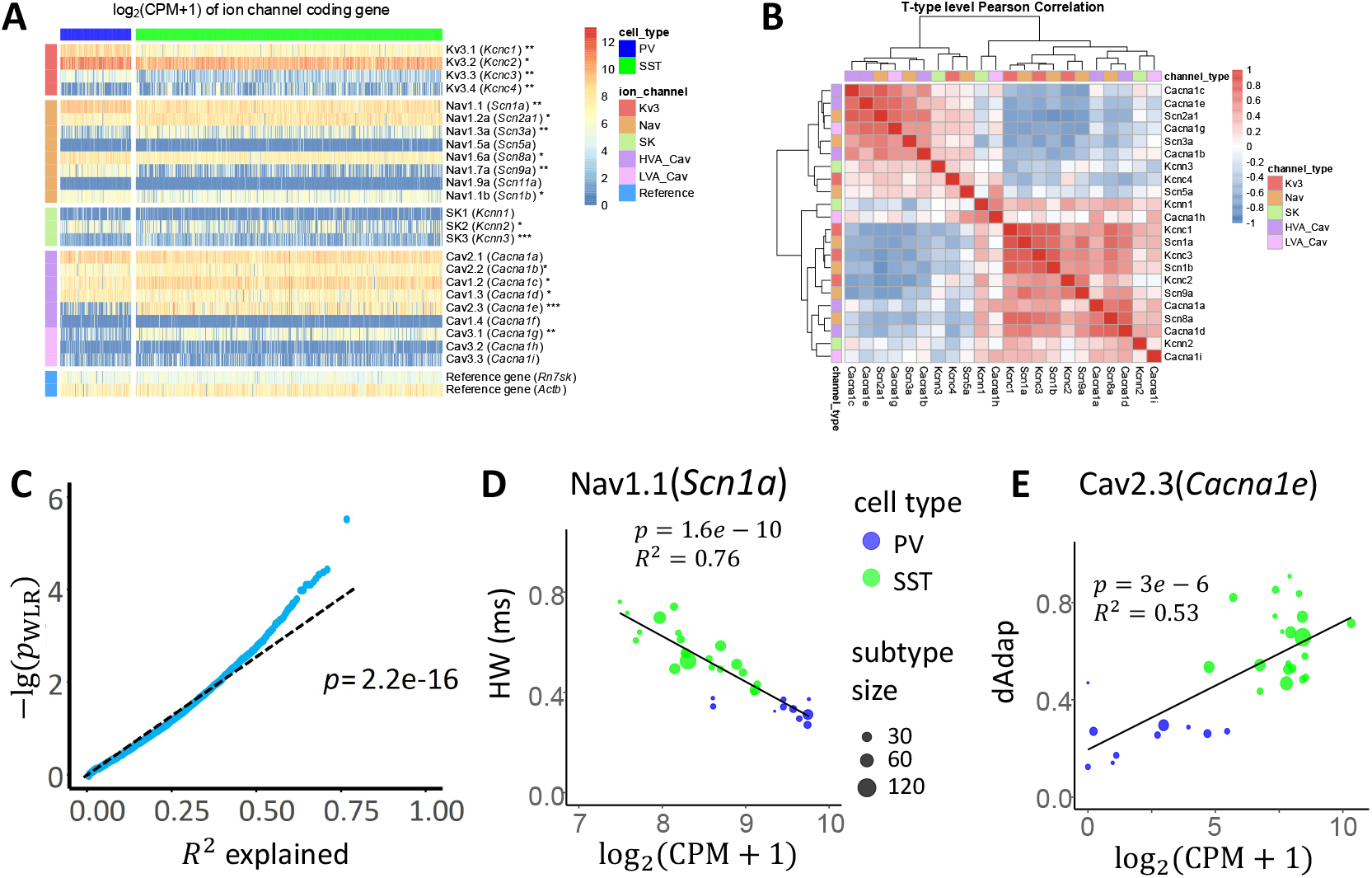
Further analyses of directly related genes. Related to Figure 3. (A) Heatmap of log_2_(CPM + 1) gene expression level at the single-cell level. Genes in rows and cells in columns. Asterisks are used to annotate differential expression significance between V1 PV and SST cells (*: fold change between 0.5 and 1, **: 1 to 2, ***: > 2). In our analysis, genes with *q* < 0.05 and fold change larger than one were counted as DEGs. (B) Person correlation of gene expression level. (C) R^2^ and −lg(*p*_*W LR*_) are linearly correlated. (D) Correlation between HW and *Scn1a*. (E) Correlation between dAdap and *Cacna1e*

**Supp. Figure S5.**
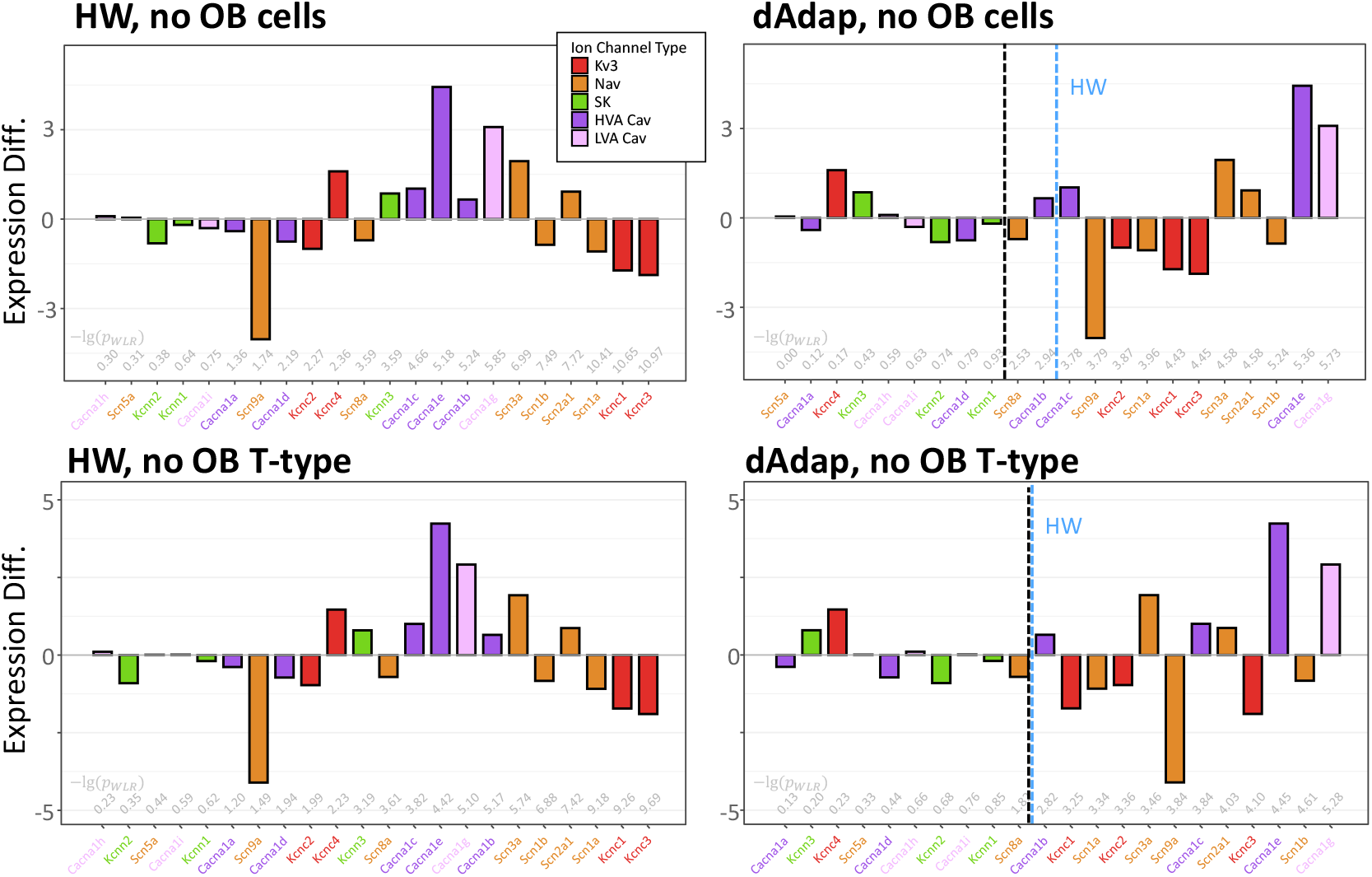
Related to Figure 3. Ranked genes that correlated with HW (Left) and dAdap (Right) in the dataset without cells with an OB feature (Top) or T-types that enriched with OB features (Bottom). The expression difference is measured by fold change of gene expressions. The black dashed line indicates the significance level *p*^*W LR*^ = 0.01. The Blue dashed line indicates −lg(*p*^*W LR*^) of HW.

**Supp. Figure S6.**
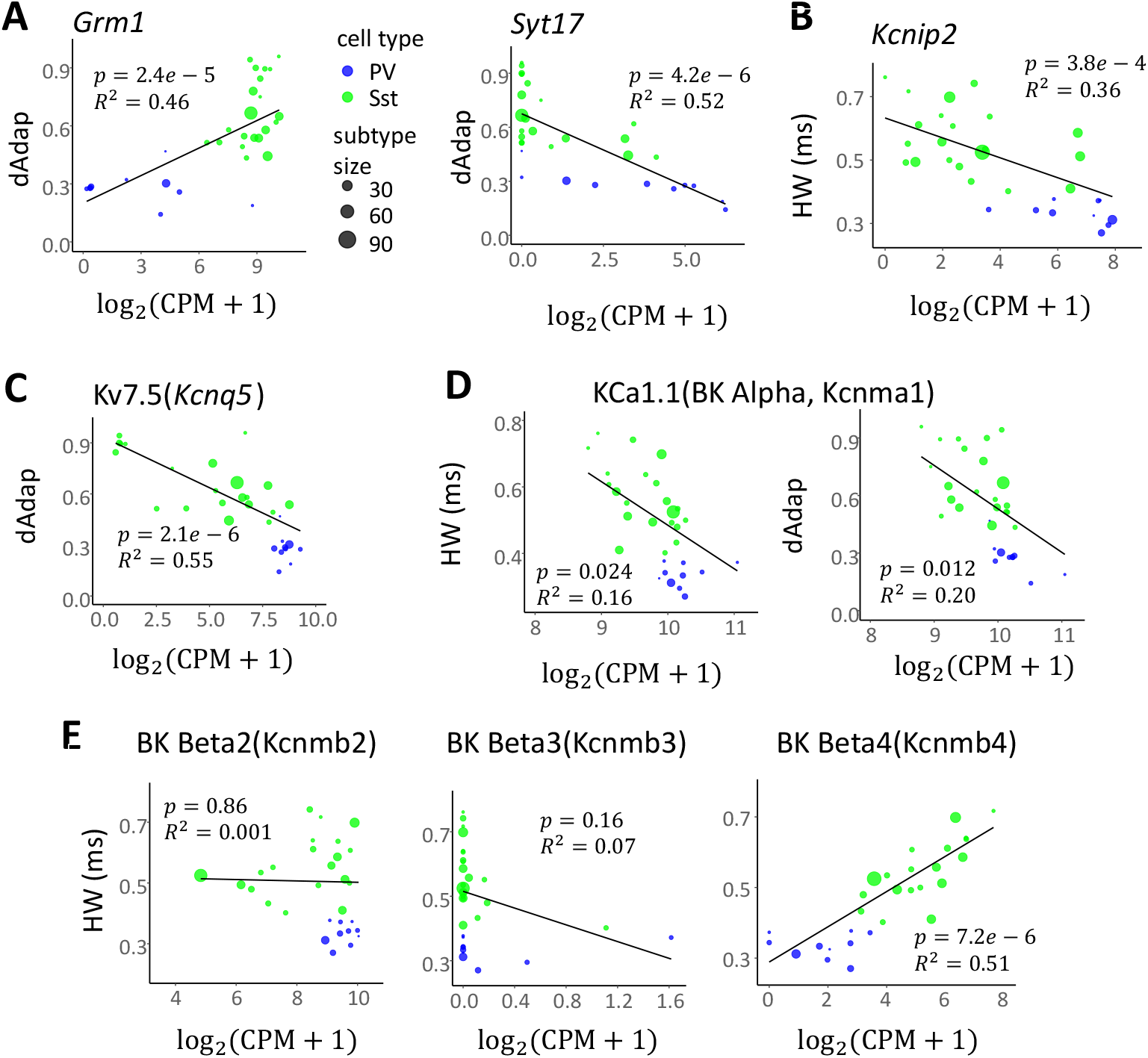
More WLR on example genes. Related to Figure 4. (A) *Grm1* and *Syt17* are highly correlated with dAdap. (B) *Kcnip2* is significantly correlated with HW. (C) *Kcnq5*, encoding voltage-gated potassium channel subunit Kv7.5, is significantly correlated with dAdap. (D) *Kcnma1*, encoding the alpha-unit of big-conductance *K*^+^ (BK) channel, are weakly correlated with HW and dAdap. (E) *Kcnmb2* to *Kcnmb4*, encoding beta-units of the BK channel, do not consistently correlate with HW.

**Supp. Figure S7.**
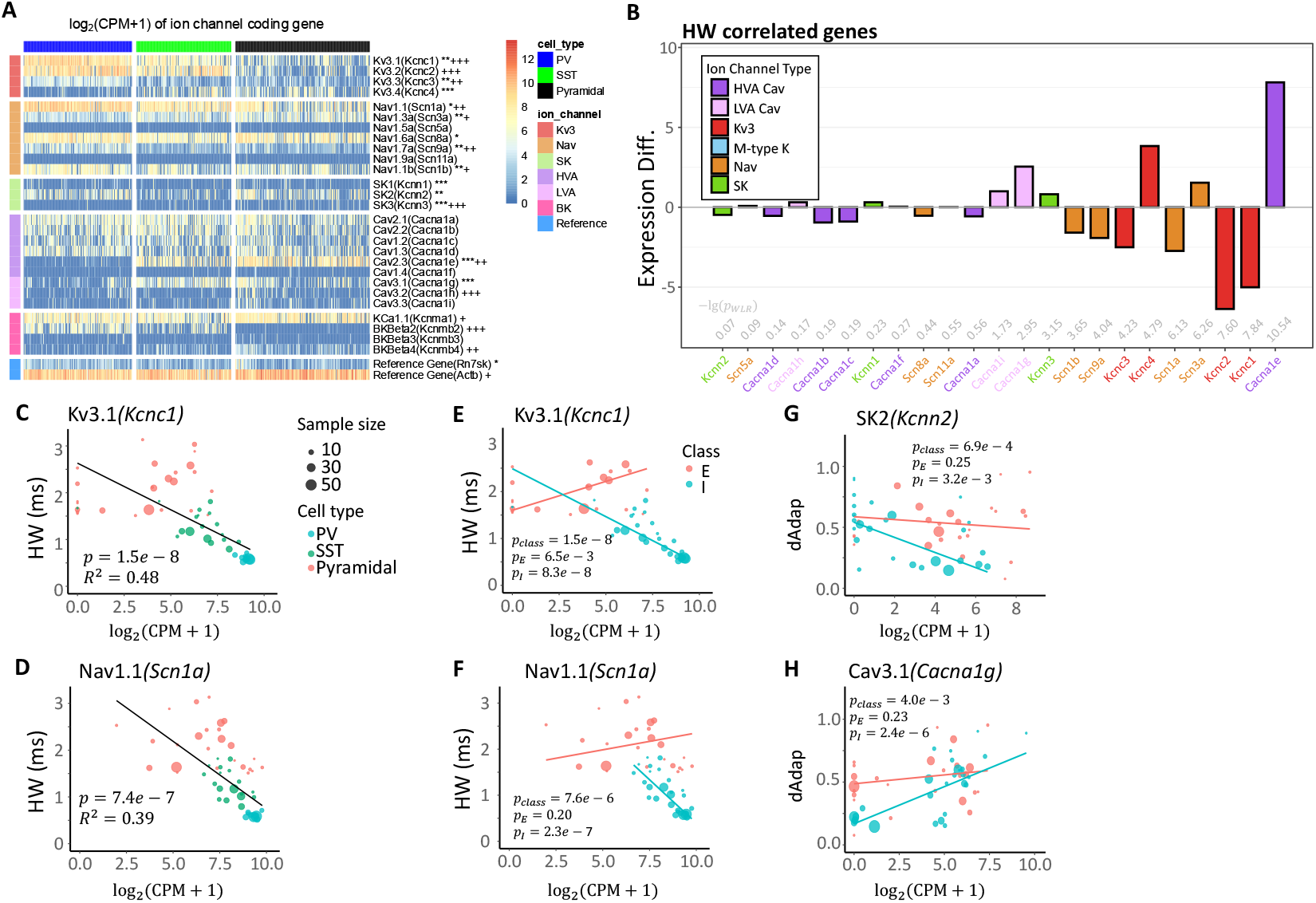
Further analyses on the mouse M1 dataset. Related to Figure 5. (A) Single-cell level heatmap and differential expression analysis. The asterisks(*) and crosses(+) represent the fold change between PV and SST, or INs and pyramidal cells, respectively (*/+: fold change between 0.5 and 1, **/++: 1 to 2, ***/+++: > 2). (B) Ranked genes that correlated with HW. The expression difference is measured by fold change. (C-D) Correlation between HW and *Kcnc1* (C) and *Scn1a* (D) across Pyramidal, SST, and PV cells. (E-H) Class-dependent effect and within-class fitting results of different genes. *p*_*class*_ shows the significance of the class-dependent effect; *p*_*E*_ and *p*_*I*_ show the significance of a non-zero slope in the I and E subsets, respectively. E and I subsets are labeled with pink and blue. See Supp. Sheet 5.

**Supp. Figure S8.**
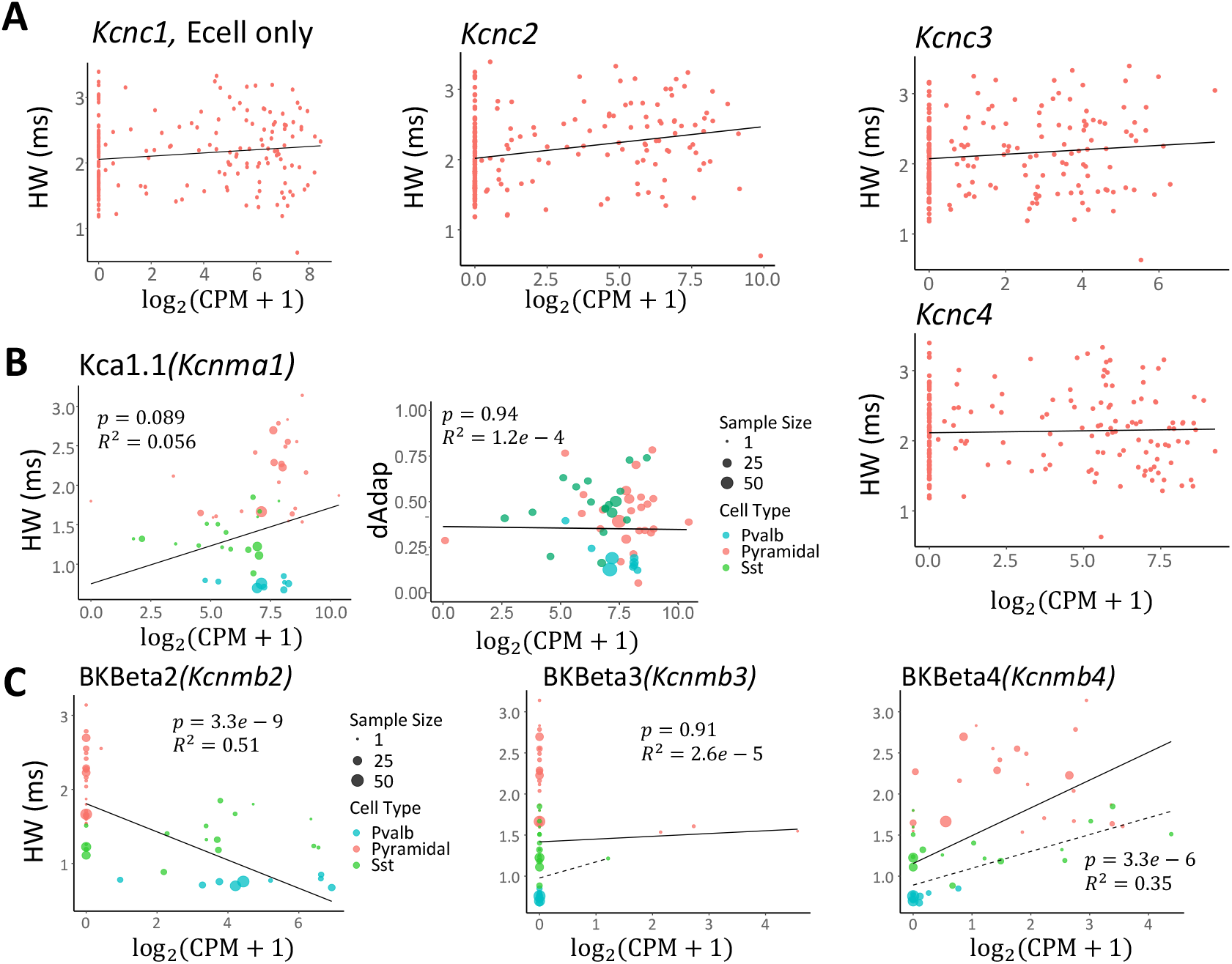
HW differences in M1 pyramidal cells can not be explained by other K channels. Related to Figure 5. (A) correlation between Kv3 encoding genes (*Kcnc1* to *Kcnc4*) and HW at the single cell level for pyramidal cells. All non-zero slopes are non-significant. (B) Correlation between BK alpha unit *Kcnma1* and HW or dAdap. (C) Correlation between BK beta unit *Kcnmb2* to *Kcnmb4* and HW. Based on (B, C), BK encoding genes cannot explain the observed HW difference within pyramidal cells.

**Supp. Figure S9.**
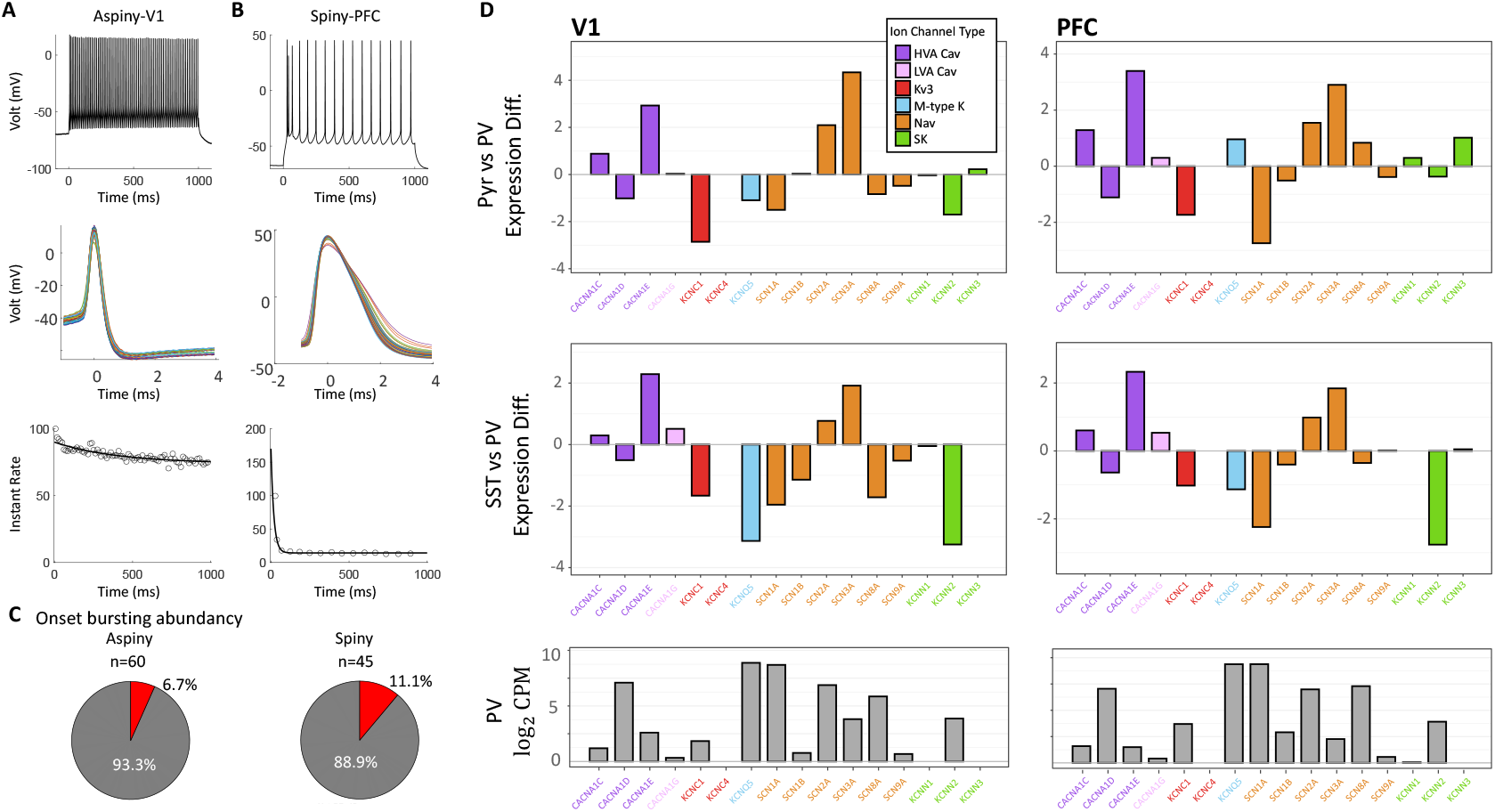
Physiology data analysis of macaque monkeys. Related to Figure 6. (A) Different electrophysiological features from a PFC spiny cell, a putative pyramidal cell. From top to bottom: recording over 1s-long square-pulse current injection; AP waveform; instantaneous firing rate as one over interspike intervals. (B) Same as (A) but from a V1 aspiny cell, a putative interneuron. (C) Abundance of OB features in aspiny cells (left) or spiny cells (right). (D) The expression difference is measured by log_2_ fold change of gene expressions in pyramidal (top) and SST cells (middle) compared to that in PV cells at V1 (left) and PFC (right). The mean expressions in PV cells are shown at the bottom. Ion channel families are color-labeled.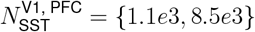

**Supp. Figure S10.**
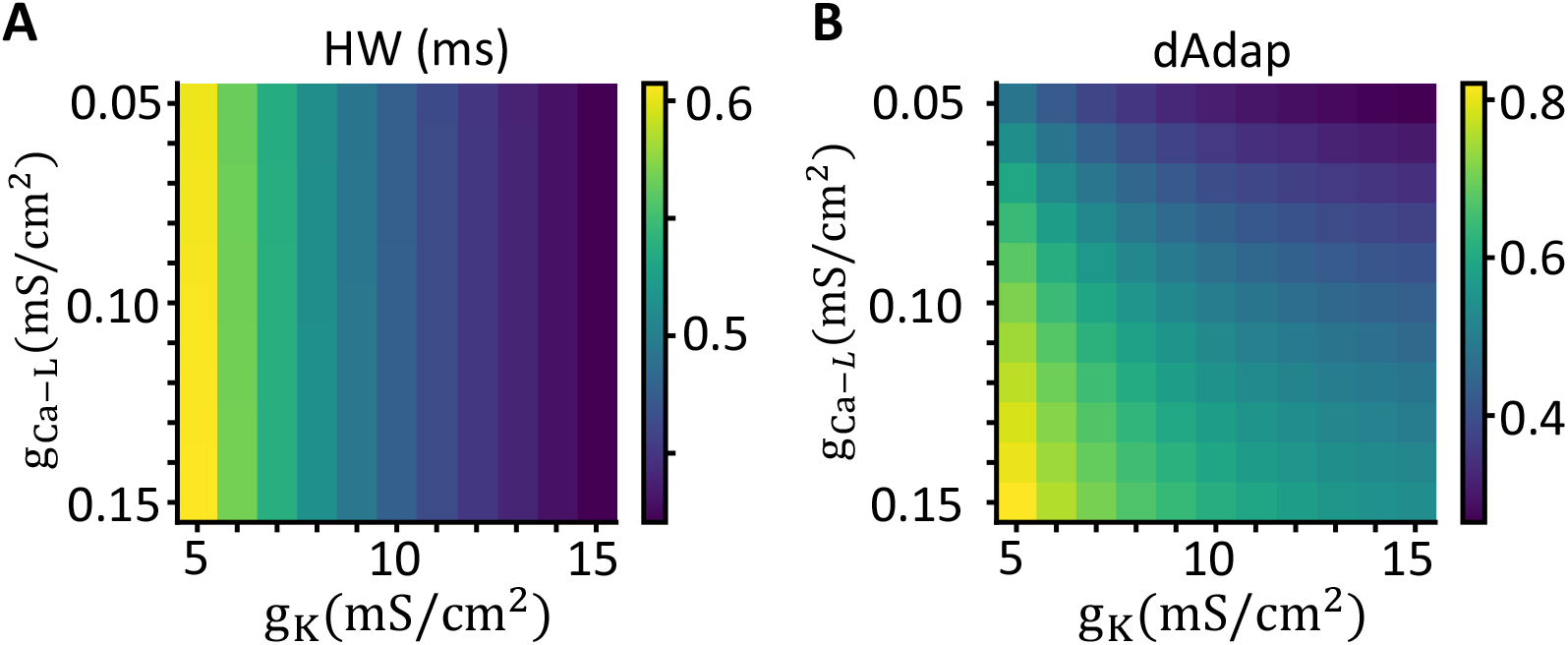
Related to Figure 7 Effects of varying *g*_*K*_ and *g*_*Ca*−*L*_ on HW (A) and dAdap (B). Reducing *g*_*Ca*−*L*_ selectively decreases dAdap without affecting HW.

